# Metabolic engineering of *Salmonella enterica* for coupled kynurenine sensing and depletion enhances antitumor efficacy

**DOI:** 10.64898/2026.09.23.753816

**Authors:** Kontham Kulangara Varsha, Akeem Santos, Tingting Xiao, Zeneng Wang, Rashmi Bharti, Mahima Arora, Austin M Williams, John Peterson, Justin D. Lathia, Stanley L. Hazen, Nan Zhang, Philip Ahern, Ofer Reizes, Mohammed Dwidar

**Author notes:** Correspondence (Mohammed Dwidar). Contributed equally as co-first authors (Kontham Kulangara Varsha, Akeem Santos).

## Abstract

Immunosuppressive tumor metabolic microenvironment remains a major barrier to effective cancer therapy. Tumor-targeting Salmonella enterica strains were engineered to deplete the intratumoral immunosuppressive kynurenine, triggering metabolic rewiring and fostering an active antitumor immune microenvironment. Equipping S. enterica VNP20009 with bacterial kynureninase enzyme (KynU) and a kynurenine transporter enabled efficient intratumoral kynurenine degradation. We then developed a next-generation strain, AD51, combining quorum-sensing controlled KynU expression with kynurenine-dependent growth and tumor targeting. AD51 induced robust immune activation and exhibited better anti-tumor efficacy. The addition of intratumoral exogenous IFNγ to AD51 synergistically enhanced immune activation and antitumor efficacy in murine ovarian cancer and melanoma models, reducing tumor burden by 75% and 84%, respectively, compared with untreated controls (p<0.0001 for both). Our data establishes a framework for engineering bacteria to target and degrade tumor immunosuppressive metabolites while coupling bacterial growth to metabolite availability, paving the way for more precise and programmable microbial therapies.

## Introduction

Kynurenine is an immunosuppressive catabolite of tryptophan overproduced by almost all solid tumors including ovarian cancer^1–3^, glioblastoma^4,5^, breast cancer^6–9^, colorectal cancer^10–12^, head and neck squamous cell carcinoma (HNSCC)^13^ and others^14^ that allows the tumor to escape immune surveillance^15,16^. The level of kynurenine production differs between tumors and is induced by the T-cell infiltration and the associated inflammatory cytokines especially IFNɣ within the tumor microenvironment^17^. The immunosuppressive and protumorgenic action of kynurenine is mainly attributed to its binding and activation of the arylhydrocarbon receptor (AhR) in the T-cells. This binding enhances conversion into Treg cells^9,18^ among other actions including recruitment of M2 macrophages and myeloid-derived suppressor cells (MDSCs) in a Treg-dependent manner^19,20^. Inhibition of indolamine 2, 3-dioxygenase 1 (IDO1), the rate-limiting enzyme in the tryptophan–kynurenine metabolic pathway, has long been proposed as a potential therapeutic strategy to enhance antitumor immune activation ^20,21^. Several IDO inhibitors have entered different phases of oncology clinical trials in combination with other immunotherapy approaches (e.g., NCT01792050, NCT02077881, and NCT02471846) with the hypothesis that countering the kynurenine pathway will lead to enhanced immune stimulation that can limit tumor growth. Nevertheless, recent clinical studies focusing on IDO1 blockade in ovarian cancer found it ineffective. Although the precise reasons remain unclear, tumor metabolic adaptation that shunted tryptophan catabolism toward the serotonin pathway may be responsible^22^. This in turn highlights the need to find alternative ways to overcome the kynurenine accumulation in tumors.

Some bacteria such as *Pseudomonas aeruginosae* harbor kynureninase (KynU) enzyme which metabolizes kynurneine into anthranilic acid which is further converted to other metabolites^23–26^. Previous studies showed that intra-tumoral administration of purified bacterial KynU enzyme successfully degraded the intratumoral kynurenine and inhibited tumor growth in melanoma, breast, and colon carcinoma models^18^. In that study, however, the authors had to PEGylate KynU to improve its stability in the tumor microenvironment^18^.

Genetically-engineered bacteria including *Salmonella enterica* and other strains are emerging as cancer therapeutics and may be a future potential treatment option for solid tumors^27–30^, and form the basis for several completed or ongoing clinical trials in which genetically modified *S. enterica* is used as monotherapy or in combination with other cancer therapies (e.g. NCT06178003, NCT05038150, NCT04589234, NCT06181266, NCT03762291, NCT03421236, NCT03750071). These bacteria both directly kill the cancer cells due to their native cytotoxicity, and more importantly, owing to their natural immunogenic properties, stimulate the immune cells within the tumor microenvironment to better attack and eradicate the tumor. Widespread application of this approach, however, is limited and there is still a need to improve the therapeutic efficacy of *S. enterica* in stimulating the immune system^31,32^.

In our previous study, we engineered *S. enterica* AD95+ strain (named hereafter as AD95) to be dependent on kynurenine for growth^33^. In murine models of breast and ovarian cancer, we demonstrated that our engineered *S. enterica* AD95 strain accumulates in tumors at levels up to 100,000-fold higher than in other organs, including the liver, spleen, kidneys, and lungs. This tumor accumulation and specificity far exceed the tumor-targeting efficiency of *S. enterica* VNP20009, the most extensively characterized tumor-specific strain in clinical trials^34–37^. Indeed, by 7-days post-intravenous bacterial injection, VNP20009 remained at high numbers in non-tumor tissues while our engineered AD95 strain almost disappeared from all other organs except tumors where they maintained robust and persistent colonization^33^.

Here, we aimed at enhancing the efficacy of the *Salmonella*-based cancer therapy in modulating the tumor immune microenvironment through engineering *S. enterica* to degrade the pro-tumorigenic immunosuppressive kynurenine secreted by the tumor cells. We first show that equipping *S. enterica* VNP20009 with kynureninase enzyme (KynU) together with kynurenine transporter enabled it to effectively degrade kynurenine in murine tumors. Furthermore, combining kynurenine targeting (parent AD95 strain) with quorum-sensing controlled KynU expression in one strain (named AD51) enabled both specific tumor-targeting and kynurenine degradation. Compared to the parent strain (AD95), which does not degrade kynurenine, AD51 exhibited better anti-tumor efficacy and enhanced immune stimulation in the tumor. Finally, intratumoral administration of IFNγ augmented the anti-tumor efficacy of AD51 in a high-kynurenine-producing KPCA.A murine ovarian cancer model as well as in a low-kynurenine-producing B16-F10 melanoma model.

## Results

### Investigating natural kynurenine metabolism in *S. enterica*

Some bacteria including *E. coli* metabolize kynurenine into kynurenic acid through transaminases such as aspartate transaminase (AspC)^38,39^. *S. enterica* ATCC 14028 naturally harbors the promiscuous aspartate transaminase (AspC) in addition to other related transaminases including aromatic amino acid transaminase (TyrB) (Figure 1A). We previously demonstrated that *S. enterica* can also metabolize kynurenine into kynurenic acid^33^. To get better insights into the native kynurenine metabolism in *S. enterica* and test if this transamination reaction in *S. enterica* is catalyzed by the AspC enzyme, we generated *S. enterica* Δ*aspC* knockout. In parallel, we also transformed the wild-type *S. enterica* with a plasmid harboring the aromatic amino acid transporter (Mtr). This transporter enhances kynurenine import^33^. Analyzing the LB culture spent media of each of these strains through LC-MS/MS confirmed that *S. enterica* naturally metabolizes kynurenine into kynurenic acid, albeit modestly (Figure 1B). The overexpression of the Mtr transporter resulted in higher kynurenic acid. Knocking out *aspC* reduced the amount of kynurenic acid produced but did not abolish it, suggesting that AspC is not the major determinant in this conversion. We anticipated this activity may be due to TyrB, and therefore, we generated and tested Δ*tyrB*, and Δ*aspC*Δ*tyrB* double knockout. The results found that knocking out *tyrB* significantly reduced the ability of *S. enterica* to metabolize kynurenine. Likewise, overexpressing *tryB* and, to a lesser extent, *aspC* significantly enhanced the natural ability of *S. enetrica* to degrade kynurenine to kynurenic acid. These results confirmed that TyrB, but not AspC is the major contributor tokynurenine metabolism into kynurenic acid (Figure 1B).

**Figure 1.**
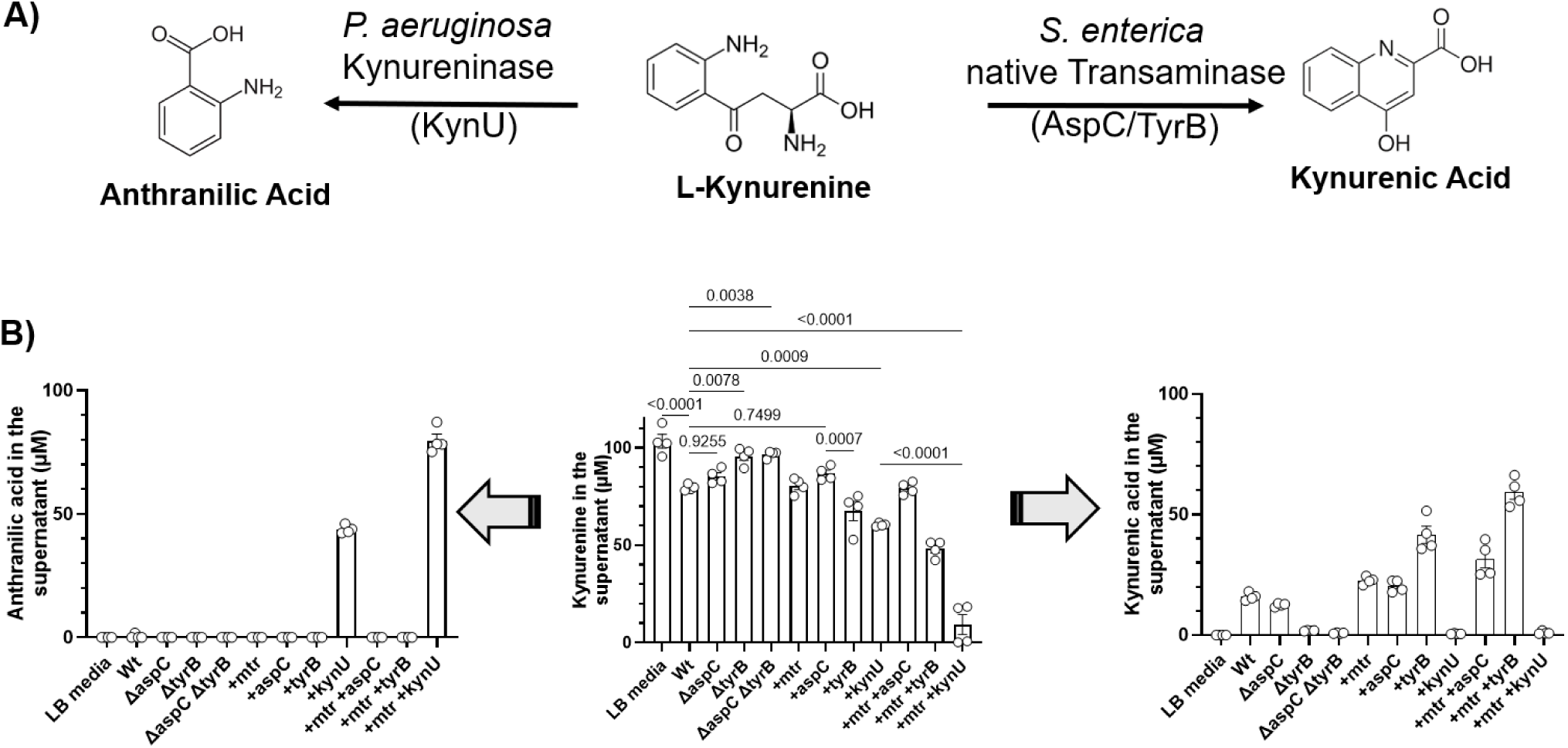
Kynurenine metabolism by wildtype *S. enterica* and mutants in LB media. **A)** Structure of kynurenine and its two metabolites; anthranilic, and kynurenic acids. **B)** Wild-type *S. enterica* (Wt) and mutants were cultured in LB media supplemented with 100 µM kynurenine. Samples were analyzed after 24 h of incubation through LC-MS/MS. n=4. Plotted are means ± SE. Statistical analyses were done using one way ANOVA.

We then investigated whether *S. enterica* could be engineered to enhance kynurenine degradation without relying on its native transaminases, which participate in central amino acid metabolic pathways and may compromise bacterial fitness. To achieve this, we explored equipping it with a recombinant kynureninase (KynU), an enzyme that converts kynurenine into anthranilic acid. Wild-type (Wt) *S. enterica* was unable to convert kynurenine to anthranilic acid as it does not harbor a KynU enzyme. However, cloning and expressing *kynU* gene from *P. aeruginosa* together with the Mtr transporter allowed *S. enterica* to degrade kynurenine to anthranilic acid and abolished its native kynurenic acid production (Figure 1B). Similar results were obtained when the experiment was repeated in defined M9 media (Figure S1A). However, in M9 media, we noticed that Wt *S. enterica* metabolized kynurenine completely to kynurenic acid in almost equimolar levels. *S. enterica* harboring KynU and Mtr, on the other hand, still metabolized kynurenine completely to anthranilic acid. However, the molar concentration of the produced anthranilic acid was almost 60% of the added kynurenine. This is likely due to the consumption of part of the produced anthranilic acid within the tryptophan biosynthesis pathway (Figure S1B). To confirm this hypothesis, Wt *S. enterica* was cultured in M9 media supplemented with either kynurenic acid or anthranilic acid. The results confirmed the ability of Wt *S. enterica* to metabolize anthranilic but not kynurenic acid (Figure S1C, D).

### Equipping *S. enterica* VNP20009 with KynU enables kynurenine degradation in murine KPCA.A tumor models

A kynurenine-degrading plasmid harboring both *mtr* and *kynU* (pMtr-KynU) was then transformed into *S. enterica* VNP200009. This strain is a derivative of the Wt *S. enterica* ATCC 14028, which is attenuated to reduce its toxicity and improve tumor targeting^34–36^. To ensure plasmid stability, transformed VNP20009 was rendered auxotrophic for D-glutamate (hereafter named VNP20009m) through knocking out the *murI* gene (needed for D-glutamate synthesis and subsequent synthesis of the peptidoglycan cell wall) ^40^. A copy of the *murI* gene was then supplied on the kynurenine-degrading plasmid, which was transformed into VNP20009m to generate the VNP20009m/ pMtr-KynU strain. Testing this mutant in LB media confirmed the enhanced capability to degrade kynurenine to anthranilic acid, similar to what we observed with the Wt *S. enterica* (Figure 2A-C).

**Figure 2.**
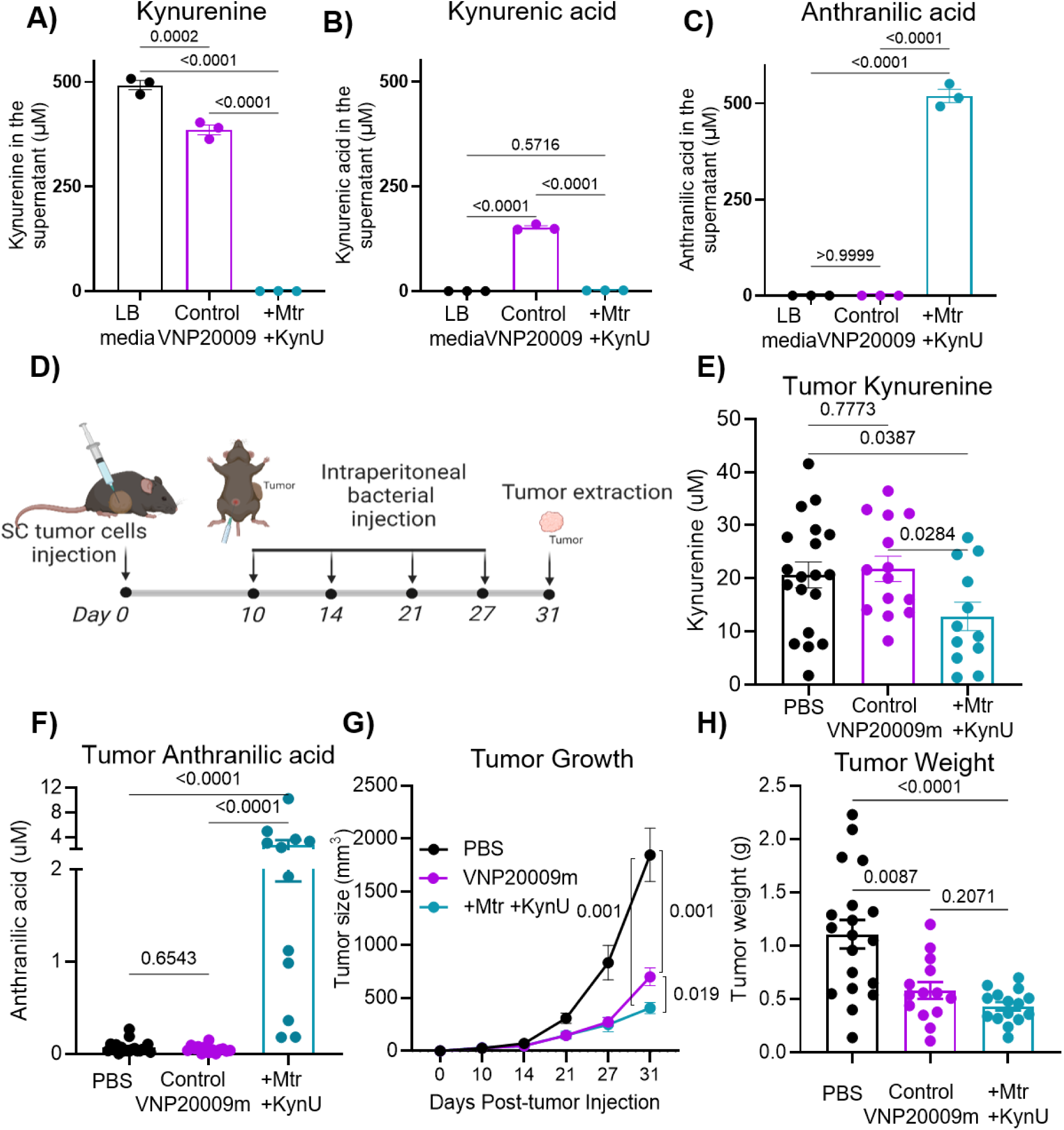
*S. enterica* VNP20009 engineered to express KynU enzyme and overexpress the Mtr transporter metabolizes kynurenine into anthranilic acid both in vitro and in murine tumors. **A-C)** *S. enterica* VNP20009 and its VNP20009m/pMtr-KynU mutant were cultured in LB media supplemented with 100 µM kynurenine. Samples were analyzed after 24 h of incubation through LC-MS/MS. n=4. **D)** Murine tumor experiment scheme. KPCA.A tumors were developed subcutaneously in C57BL/6 mice. When the tumors were evident, VNP20009 mutants were intraperitoneally injected at a dose of ∼ 2×10^6^ CFU weekly. The mice were euthanized 31 days after the tumor injection (when the tumors in the PBS group reached the endpoint). n=7∼10 per group. Tumor size was measured weekly while tumor weight was determined at the endpoint. **E, F)** Tumor kynurenine and anthranilic acid concentrations. **G)** Tumor growth over time. **H)** Tumor weight at endpoint. For panels E-F, data were combined from two separate experiments. Statistical analyses were done using one way ANOVA for A-C and Kruskal-Wallis test for E-H. Plotted are means ± SE.

To test the ability of VNP20009m/ pMtr-KynU strain to degrade tumor kynurenine to anthranilic acid in vivo, we established subcutaneous tumors in C57BL/6J mice using the KPCA.A murine ovarian cancer model^41^. The mice were i.p. injected weekly with either PBS, Control VNP20009m harboring only the MurI encoding plasmid or the VNP20009m/ pMtr-KynU mutant (Figure 2D). The results confirmed the capability of KynU-harboring *S. enterica* to degrade tumor kynurenine into anthranilic acid (Figure 2E, F). Further, the reduction in tumor growth was more evident in the VNP20009/pMtr-kynU group compared to the parent VNP20009m treated-mice suggesting a therapeutic benefit of the KynU-expressing *S. enterica* treatment (Figure 2G, H). No difference in tumor colonization and specifity were seen between the two strains (Figure S2).

### Combining kynurenine targeting and degradation in the same *S. enterica* strain

In our previous study, we developed a tumor-targeting *S. enterica* AD95 strain. In contrast to VNP20009, which is auxotrophic for purines, AD95 is dependent for its growth on kynurenine^33^. This dependency was attained through deleting the *murI* and *asd* genes needed for the synthesis of D-glutamate and diaminopimelic acid (DAP), respectively. These two genes were then supplied *in trans* under kynurenine-controlled genetic circuits (Figure 3A)^33^. We demonstrated that AD95 accumulates in tumors at levels up to 100,000-fold higher than in other organs, including the liver, spleen, kidneys, and lungs. This tumor accumulation and specificity far exceed the tumor-targeting efficiency of *S. enterica* VNP20009^33^. We therefore aimed at moving our kynurenine degradation modules to the more tumor-specific AD95 strain. For our first engineered AD95 derivative (hereafter named AD31), KynU enzyme was expressed in the chromosome downstream of the constitutive synthetic promoter PJ23114 (Figure 3B). Unlike the parent AD95, AD31 was unable to grow in M9 defined media in the absence of DAP and D-glutamate, even when supplemented with kynurenine (Figure 3D-F). That was expected since both AD95 and AD31 need kynurenine as an inducer for growth. Forcing AD31, therefore, to metabolize kynurenine constitutively would limit the kynurenine available to induce the expression of the two essential genes *murI* and *asd*.

**Figure 3.**
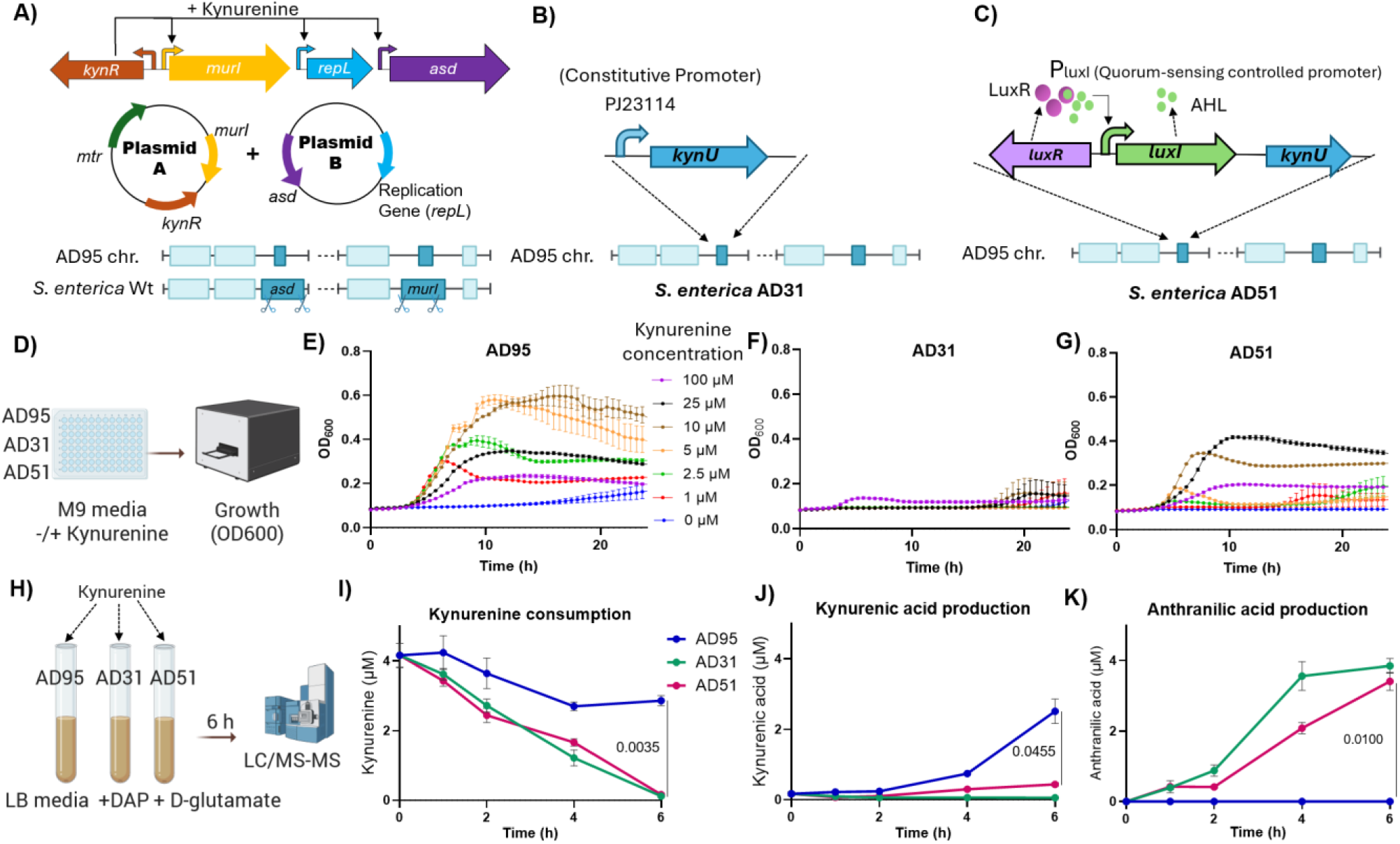
Engineering AD31 and AD51 strains from the parent *S. enterica* AD95 strain. **A)** Schematic diagram for the AD95 strain as described previously^33^. AD95 was engineered to be dependent on kynurenine for growth through knocking out *murI* and *asd* genes in *S. enterica* ATCC14028s and supplying copies of these two genes on plasmids downstream of kynurenine-responsive promoters. The replication protein of the *asd*-harboring plasmid was also placed downstream of another kynurenine-responsive promoter to further tune the growth in response to kynurenine. **B)** AD31 strain was made through knocking in the kynU gene in the chromosome of AD95 downstream a constitutive promoter (PJ23114)**. C)** AD51 was developed similar to AD31, however, the PJ23114 promoter was replaced with the LuxR-P_LuxI_-LuxI cassette so that both LuxI and KynU expression is driven by P_LuxI_ Promotor. **D-G)** Growth of each of the three strains in M9 media at different kynurenine concentrations. M9 media was supplemented with glucose at 0.4% and casamino acids at 1%. **H-K)** Kinetics of kynurenine degradation by each of the three strains in LB media supplemented with DAP and D-glutamate. These two metabolites were added to exclude the dependance on kynurenine for growth. n=3.

Consequently, we sought to control *kynU* gene expression through Acyl homoserine lactone (AHL)–based quorum sensing (Figure 3C). AHL-quorum sensing (QS) in its native bacterium, *Vibrio fischeri,* controls the transcription of the luminescence (lux) operon^42,43^. In this system, the *luxI* gene codes for AHL synthase, while LuxR acts as a transcriptional regulator which, upon binding to the autoinducer AHL, activates transcription from the promoter upstream of the *lux* operon, leading to upregulation of the *luxI* gene together with the other *lux* genes downstream of *luxI*. When the bacteria are present in low numbers, the concentration of the produced AHL molecules is very low. However, when the bacteria grow to sufficient numbers, the concentration of the produced AHL in the medium reaches a critical threshold, which allows it to bind to LuxR and activate the expression of the *lux* operon in a positive feedback loop. Expressing *kynU* under the control of the LuxI-LuxR QS system will therefore ensure that KynU is produced only after bacterial accumulation in the tumor to sufficient biomass. Indeed, when we replaced PJ23114 with the LuxR-PLuxI cassette, the resultant strain (hereafter named AD51) could grow in M9 media without DAP or D-glutamate when supplemented with kynurenine in a pattern akin to the parent AD95 strain (Figure 3G).

We next examined the capability of each of the three strains (AD95, AD31, and AD51) to degrade kynurenine in the media. To exclude the dependence on kynurenine for growth, the experiment was done in rich LB media supplemented with both DAP and D-glutamate. The results confirmed the ability of both AD31 and AD51 to degrade kynurenine almost completely into anthranilic acid (Figure 3H-K). Similar to the Wt *S. enterica* in Figure 1, AD95 converted only a fraction of kynurenine into kynurenic acid, which was not metabolized further.

### *S. enterica* AD51 successfully targets and degrades tumor kynurenine in vivo

To assess the ability of AD51 to retain tumor specificity, we established KPCA.A subcutaneous tumors in C57BL/6J mice as above. When the tumors reached medium size, the mice were intravenously injected with either the parent kynurenine-targeting AD95 strain or the modified kynurenine-targeting and degrading AD51 strain. One week later, the mice were euthanized, and bacterial counts in tumors and other organs were determined through plating. We found that AD51 retains high specificity to the tumor akin to the parent AD95 strain (Figure S3). Surprisingly, there was a slight enhancement in tumor colonization compared to AD95. However, colonization of other organs was also slightly increased, resulting in similar tumor/liver and tumor/spleen ratios as AD95 (Figure S3C-D). This slight enhancement in colonization might be due to the ability of AD51 to use kynurenine as a carbon source rather than only as an inducer in the case of AD95. It should be noted, however, that this phenomenon was not observed for VNP20009 when supplemented with kynurenine-degrading ability (Figure S2).

We next opted to assess the in vivo kynurenine-degrading and tumor-mitigating effects of AD51 compared to AD95. KPCA.A tumors were established subcutaneously as explained above. The bacteria or control PBS solution was injected intravenously after the tumors were evident. Tumor growth was assessed weekly, and all mice were euthanized when tumors in the PBS group reached the endpoint. The results showed that both strains significantly reduced tumor growth over time and tumor weight at the endpoint (Figures 4A-C, S4A). AD51, however, was more effective in reducing tumor burden. This was consistent with the reduction in tumor kynurenine seen in the AD51 group but not in the AD95 group (Figure 4D-E). To explore how kynurenine degradation, when combined with *S. enterica* treatment, affected the tumor immune microenvironment, the tumors in each group were analyzed for their cytokine content using multiplex ELISA. As expected, the tumors treated with bacteria (either AD95 or AD51) showed elevated levels of multiple proinflammatory cytokines, which are typically raised in infections, including TNFα, IL-1β, KC/GRO, and IL-6 (Figure 4F-I). While IL-5, IL-10, IL-4, and IL-12p70 did not show difference compared to the PBS group (Figure 4J-M). It was interesting, however, to see that IFNγ and IL-2 were only significantly elevated in the AD51 group but not in the AD95 group (Figure 4N-O), suggesting the polarization of a an immune signature that tracked with enhanced capacity to limit tumor burden.

**Figure 4.**
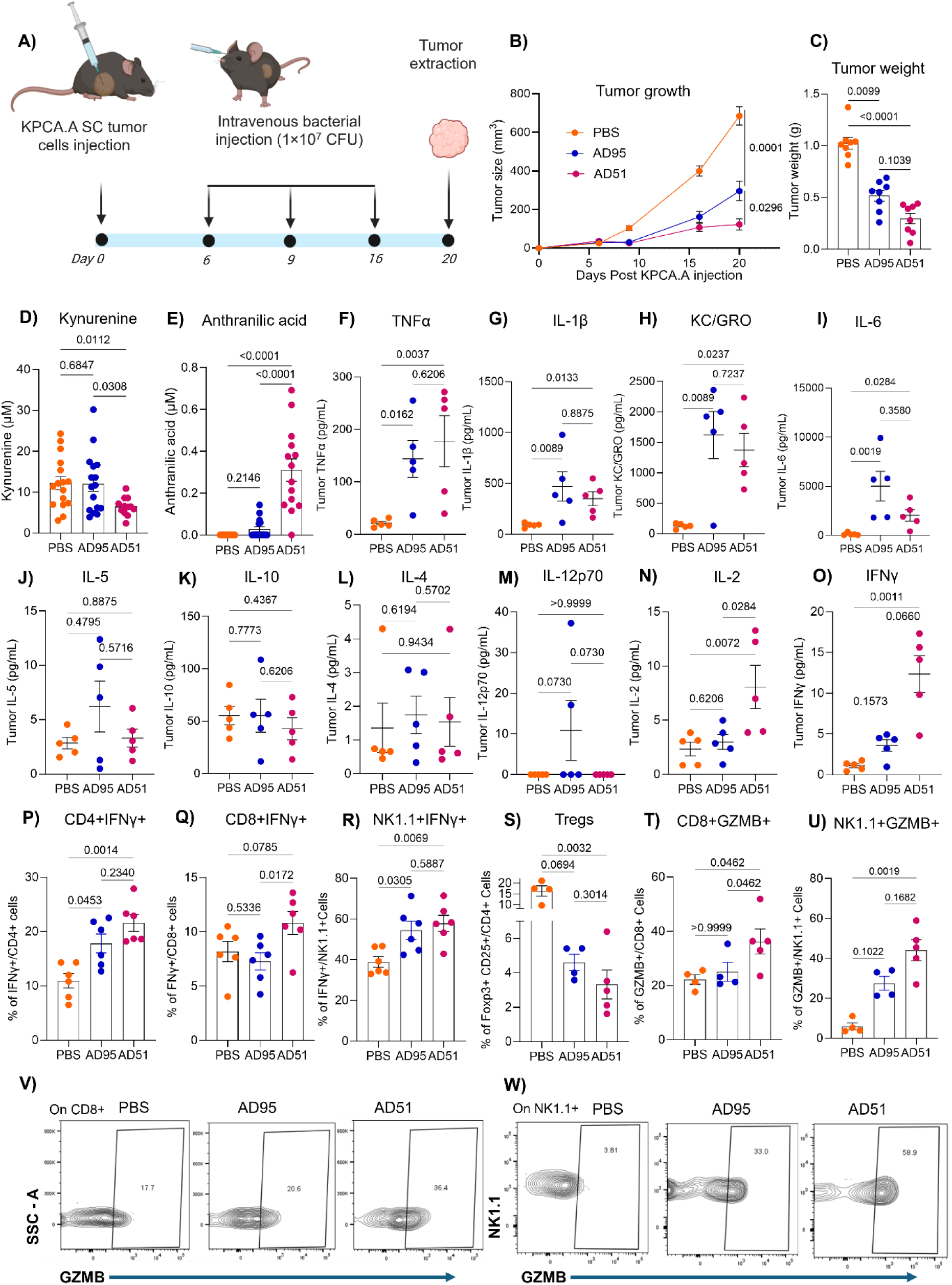
Tumor suppression by the engineered AD51 is associated with tumor immune stimulation. **A)** Experiment scheme. KPCA.A tumors were developed subcutaneously in C57BL/6 mice. When the tumors were palpable, *S. enterica* mutants were intravenously injected at a dose of ∼ 1×10^7^ CFU as mentioned in the experiment scheme. n=8∼9 per group. Tumor size was measured weekly while tumor weight was determined at the endpoint. **B)** Tumor growth over time. **C)** Tumor weight at endpoint. **D, E)** Tumor kynurenine and anthranilic acid concentrations. For panels **D** and **E**, data were combined from two separate experiments. **F-O)** Cytokine analysis by multiplex ELISA. **P-W)** Flow cytometry analysis for tumor immune cells. **P-R)** IFNγ production by CD4+ T cells, CD8+ T cells and NK cells. **S)** CD4+ CD25+ FOXP3+ Treg cells. **T, V)** Granzyme B production by CD8+ T cells. **U, W)** Granzyme B production by NK cells. Statistical analyses were done using Kruskal-Wallis test. Plotted are means ± SE.

We next performed flow cytometry-based phenotyping of tumor-residing immune cells following *S. enterica* treatment (Figure 4P-W, S4C-O, and S5). We found that the production of IFNγ by CD8+ T cells was markedly increased in the AD51 treatment group compared to the control and AD95 treatment groups. IFNγ production by CD4+ T and NK cells also showed an increase in both bacteria-treated groups compared to untreated ones (Figure 4P-R), while a significant increase in TNFα production was observed only by CD4+ T cells in both treatment groups compared to the untreated control group (Figure S4 I-N). A decrease in the frequency of CD4+CD25+Foxp3+ Tregs was observed in both bacteria-treated groups, with a more evident reduction for AD51 treatment (Figure 4S, S4O). Furthermore, we found AD51 treatment markedly increased the population of granzyme B (GZMB)+ cytotoxic CD8+ T cells (Figure 4T, V) and GZMB+ cytotoxic NK cells (Figure 4U, W). Overall, compared with the parental AD95 strain, treatment with AD51 resulted in greater activation of T cells and NK cells, along with a reduced abundance of Tregs, suggesting it mediates its superior anti-tumor effects through immune activation.

### Intratumoral IFNγ administration further augments the tumor-suppressive efficacy of *S. enterica* AD51

Recent studies found that combining IFNγ with microbial products, which serve as pattern recognition receptor (PRR) agonists, is a promising strategy to treat tumors, including ovarian cancer. These microbial products included β-glucan^44^ in one study on ovarian cancer and monophosphoryl lipid A (a derivative of bacterial LPS) in another study by a different group focusing on breast cancer^45^. Although the mechanisms are still not fully understood, these studies support the idea that microbial products synergize with IFNγ to activate a strong immune response against tumors. At the same time, IFNγ is the main inducer of kynurenine production in the cancer cells^17^. Indeed, our previous results found IFNγ addition to KPCA.A and other ovarian and breast cancer cell lines led to a large production of kynurenine in in vitro tissue culture experiments^33^. We therefore asked if the provision of external IFNγ along with our engineered *S. enterica* AD51 would further enhance the impact of AD51, hypothesizing that IFNγ would lead to more kynurenine production by the cancer cells, which would in turn enhance the tumor-localized growth of AD51, leading to additional tumor regression. In return, AD51 could enhance the immune-stimulatory effects of IFNγ. To test this hypothesis, KPCA.A tumors were generated as mentioned above. When tumors were evident, bacteria (AD95 and AD51) were injected i.v. while IFNγ was injected intratumorally as outlined in Figure 5A. Monitoring the tumor growth over time and tumor weight at the endpoint found that IFNγ injection alone did not reduce tumor growth in the PBS group mice. When IFNγ was administered to the AD51-treated tumors, a significant reduction was observed compared to injecting IFNγ or AD51 alone. A slight therapeutic advantage was also observed when IFNγ was administered to the AD95-treated group when compared to AD95 treatment alone, but the effect was not statistically significant and less prominent when compared to AD51+IFNγ treatment (Figure 5B-C). Interestingly, IFNγ did not significantly affect intratumoral AD95 or AD51 colonization levels (Figure 5D). As expected, treatment with AD51 reduced the tumor kynurenine concentration by transforming it into anthranilic acid (Figure S6A, B).

**Figure 5.**
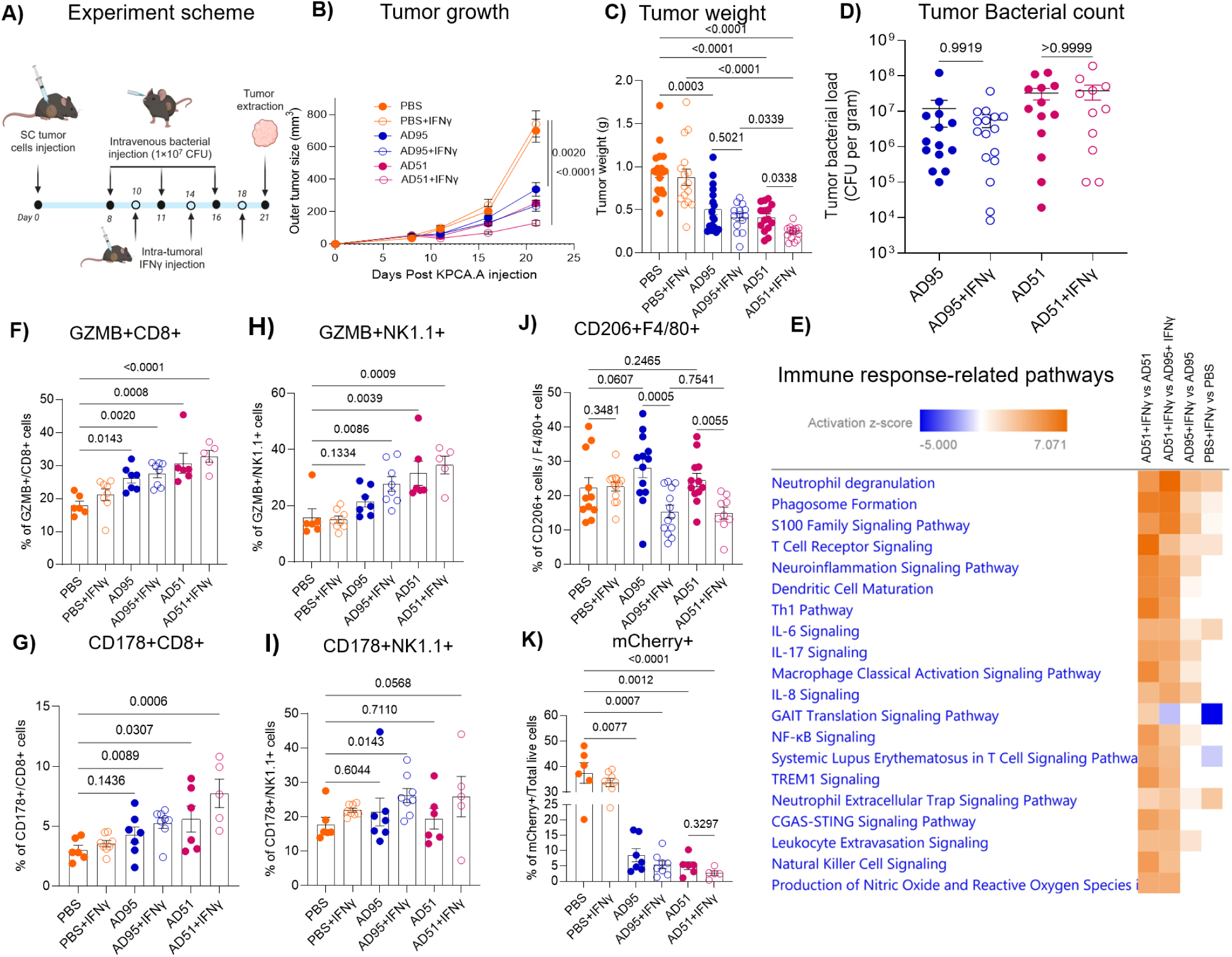
Intratumoral IFNγ administration augments the tumor-suppressive efficacy of AD51. **A)** Experiment scheme. KPCA.A tumors were developed subcutaneously in C57BL/6 mice. When the tumors were evident, *S. enterica* mutants were injected intravenously and 5 µg of IFNγ was injected intratumorally. The mice were euthanized 20 days after the tumor injection. n=8∼9 per group. Tumor size was measured weekly while tumor weight was determined at the endpoint. **B)** Tumor growth over time. **C)** Tumor weight at endpoint. **D)** Tumor bacterial count at the endpoint. **E)** Top 20 immune response related pathways which showed differential expression upon IFNγ addition to the treatment. The z-score predicts the activity state of the different pathways. An orange bar indicates positive z-score (Upregulation) while blue bar indicates negative z-score (Downregulation). **F-K)** Flow cytometry analyses of the tumor showing **F, G**) Cytotoxic CD8+ T cells, **H, I**) Cytotoxic NK cells, **J**) CD206-expressing F4/80+ macrophages and **K**) mCherry+ KPCA.A tumor cells. Statistical analyses were done using Kruskal-Wallis test. Plotted are means ± SE.

We next investigated how IFNγ treatment improved the antitumor efficacy of AD51. For this, we performed transcriptome analysis to find out the differences in gene expression between the groups. Pathway enrichment analysis revealed activation of a wide range of pathways associated with immune response when IFNγ was introduced along with either of the two *S. enterica* strains (Figure 5E). AD51+IFNγ treatment showed the greatest efficiency to induce robust immune response within the tumor when compared to other IFNγ treatment groups, including AD95+IFNγ treatment (Figure 5E). A list of all pathways differentially expressed for AD51+IFNγ group when compared to AD51 group with p values and z-score is provided in Table S1. The Top 15 pathways altered when IFNγ was injected along with AD51 (AD51+IFNγ vs AD51) included upregulation of Th1 pathway (z-scores 6.1), natural killer cell signaling (z-score 4.7), and macrophage classical activation (z-score 5.4) (Figure S6C-F). Several other immune-related pathways were also upregulated in the AD51+IFNγ group compared to the AD51 group, including, for example, the antigen presentation pathway (z-score 2.8) and CD40 signaling (z-score 2.1) (Table S1, Figure S6G-H).

To further confirm the immunostimulatory effects of the combined treatment (AD51 + IFNγ), we performed flow cytometry analysis of dissociated tumors from each group. We observed increased expression of CD178 (FasL) and GZMB in both CD8+ T cells and NK cells (Figure 5F–I, S7). In addition, CD206, an M2 macrophage marker, was significantly decreased when IFNγ was used in combination therapy (Figure 5J). As expected, the number of live KPCA.A cancer cells decreased following treatment with either AD95 or AD51, with an even greater reduction upon addition of IFNγ (Figure 5K).

### AD51+IFNγ combination therapy suppresses the growth of low-kynurenine producing B16-F10 tumor

We next asked if this dual treatment (AD51+IFNγ) will also provide therapeutic advantage in other tumor models which do not produce high levels of kynurenine. To answer this question, we established subcutaneous B16-F10 tumors. In contrast to KPCA.A, B16-F10 melanoma tumors do not naturally accumulate high levels of kynurenine ^20^. When B16-F10 tumors were palpable, themice were treated with AD95 and AD51 with/ without intratumoral IFNγ administration as done with the KPCA.A model (Figure 6A). Similar to what was observed with KPCA.A tumors, IFNγ treatment did not significantly affect either AD95 or AD51 colonization within the tumors (Figure 6B). Monitoring tumor growth over time and tumor weight at the end revealed that AD51+IFNγ treatment resulted in the most significant reduction in tumor growth (Figure 6C-D). Analyzing the tumor kynurenine levels through LC-MS/MS confirmed that untreated control B16-F10 tumors do not naturally accumulate high levels of kynurenine (Figure 6E). Treatment with IFNγ or AD95 but not with AD51, however, significantly elevated the kynurenine levels, while AD51 but not AD95 treatment elevated the anthranilic acid concentrations in the tumor (Figure 6F). Cytokine analysis of the tumors was performed by multiplex ELISA. Consistent with the results of KPCA.A cytokine analysis, the tumors treated with bacteria (either AD95 or AD51) showed elevated levels of TNFα, IL-1β, KC/GRO, and IL-6 compared to the untreated PBS group (Figure 6F-I), while the elevation in IFNγ and IL-2 reached statistical significance for the AD51 but not for the AD95 group (Figure 6N-O). At the same time, IFNγ administration significantly increased the amounts of IL-5, IL-12p70, IL-10 and IL-2 in AD51+IFNγ group (Figure 6J-K, N).

**Figure 6.**
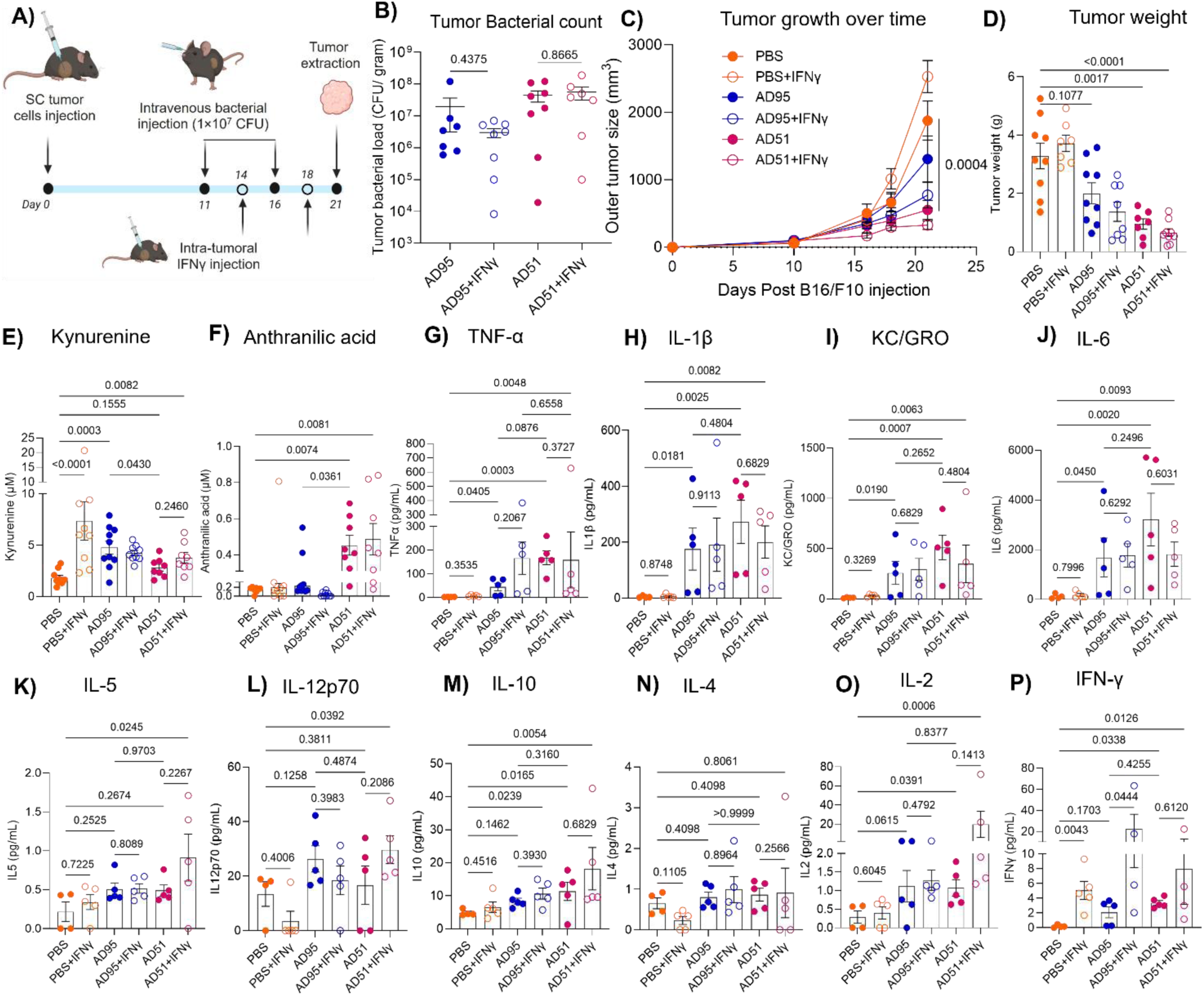
B16-F10 Tumor suppression by the engineered *S. enterica* AD95 and AD51 in combination with intratumoral IFNγ administration. **A)** Experiment scheme. B16-F10 tumors were developed subcutaneously in C57BL/6 mice. When the tumors were evident, *S. enterica* mutants were injected at a dose of ∼ 1×10^7^ CFU intravenously and 5 µg of IFNγ was injected intratumorally as mentioned in the experiment scheme. n=8∼9 per group. Tumor size was measured weekly while tumor weight was determined at the endpoint. **B)** Tumor bacterial count. **C)** Tumor growth over time. **D)**Tumor weight at endpoint. **E)** Tumor kynurenine. **F)** Tumor Anthranilic acid. **G-P)** Cytokine analysis by multiplex ELISA. Statistical analyses were done using Kruskal-Wallis test. Plotted are means ± SE.

## Discussion

Bacterial-based cancer therapy is an emerging approach which has a high potential to revolutionize the current standard of care for solid tumors at advanced stages. However, the natural immunostimulatory characteristics of the bacteria alone may not be sufficient, and novel ways to enhance the immune-stimulatory efficacy of these bacterial therapeutics are needed. One approach to achieve this is through engineering the bacteria to reshape the tumor metabolism in a way that will improve immune surveillance. Kynurenine is a key immunosuppressive metabolite in many solid malignancies ^1–14^, yet therapeutic efforts to block its production through IDO1 inhibition have not been successful so far in the clinic ^22,46^. In this study, we demonstrate that depleting kynurenine within the tumor microenvironment through the bacterial KynU enzyme effectively reprograms antitumor immunity when mediated by a tumor-targeting *Salmonella enterica*. By converting kynurenine into anthranilic acid, we abolished kynurenic acid production from the engineered *Salmonella* (Figures 1B and 3J). This observation is of particular importance in light of previous studies implicating kynurenic acid in protumorigenic and immunosuppressive pathways^47^. Importantly, we achieved this not only in the widely used VNP20009 strain but also in *S. enterica* AD95^33^, a chassis which depends on kynurenine for growth (*i.e.* AD51) without compromising the bacterial fitness and growth within the tumors, and without interfering with the selective tumor-homing of the therapeutic bacteria. By placing KynU enzyme under a fine-tuned quorum-sensing genetic circuit, we ensured that kynurenine degradation occurred only after *Salmonella* accumulates to sufficient bacterial biomass within the tumor. This design preserves our initial intended role of kynurenine as a tumor-specific growth signal during early colonization while enabling its subsequent sustained elimination once bacteria accumulate in the tumor to a critical density. Thus, AD51 functions as a self-regulating system that couples tumor targeting, metabolic intervention, and immune modulation within a single microbial chassis. This provides the added advantage of limiting interference with kynurenine’s role at non-tumor sites. Mechanistically, we found *Salmonella* colonization within the tumor when coupled with kynurenine degradation profoundly alters the tumor immune microenvironment. While both the parent AD95 strain (which does not degrade kynurenine) and AD51 (which degrades kynurenine) induced inflammatory cytokines typical of bacterial infection, AD51 treatment resulted in a more robust elevation of IFNγ and IL-2, accompanied by enhanced CD8⁺ T-cell and NK-cell cytotoxic activity.

IFNγ is known for its anti-tumorgenic effects^48^. These therapeutic effects were found in previous studies to be highly augmented when combined with microbial products which act as PRR ligands^44,45^. However, one drawback for IFNγ therapy is that it induces the tumor cells to produce immunosuppressive kynurenine (Figure 6E). With AD51 in hand, we envisioned that this drawback could be overcome. Our data demonstrates that IFNγ in combination with *S. enterica* becomes significantly more beneficial when *S. enterica* is engineered to actively deplete kynurenine from the system (Figures 5 and 6). In AD51-treated tumors, IFNγ production was amplified, creating a positive feedback loop in which immune activation fuels bacterial growth through enhanced kynurenine production, while bacterial degradation of kynurenine prevents accumulation of this immunosuppressive metabolite. It has to be noted, however, that we have not observed a significant increase in the AD51 tumor colonization when additional IFNγ was administered suggesting there might be a limit for the bacterial density within the tumors (Figures 5D and 6B).

Importantly, the therapeutic benefit of AD51 in combination with IFNγ extended beyond kynurenine-rich KPCA.A tumors. In the B16-F10 melanoma model, which exhibits low basal kynurenine levels^20^, IFNγ administration induced kynurenine accumulation that was efficiently countered by AD51. The AD51+ IFNγ combination resulted again in superior tumor control compared to treatment with AD51 or IFNγ alone, demonstrating that this dual treatment platform can function across tumors with variable kynurenine levels.

From a translational perspective, this work establishes several advances relevant to the clinical development of bacterial-based cancer therapies. First, kynurenine depletion represents a needed step toward overcoming tumor-induced immune suppression. Second, coupling tumor-specific growth of the therapeutic bacteria to reshape the tumor metabolic microenvironment enables better spatial control of the therapeutic strategy, limiting unwanted off-target effects. Finally, the rational integration of IFNγ signaling with microbe-mediated immune stimulation yields synergistic effects that exceed those of either modality alone. Compared with existing tumor-targeting *Salmonella* strains such as VNP20009, AD51 offers improved specificity (Figures S2 and S3D) and a clearly defined mechanistic basis for immune activation.

One limitation of our study, however, is that IFNγ administration is not a clinically favored modality because of its wide range of side effects^49^, and its intratumoral administration, as used here, is not always practical in clinical settings. In addition, we still need to better understand the main mechanisms by which IFNγ synergizes with the microbial therapeutic to elicit the observed antitumor response.

Nevertheless, our study demonstrates that programmable anticancer bacteria can be engineered to function as metabolite-responsive targeted agents while also metabolizing these cancer-specific metabolites within the tumor microenvironment to elicit an additional therapeutic benefit. Using the immunosuppressive metabolite kynurenine as an example, this strategy may accelerate the clinical use of bacterial-based cancer therapy and reveal new ways to reshape tumor metabolism to enhance immunity.

## Materials and Methods

### Bacterial strains and plasmids

*S. enterica* ATCC 14028, *S. enterica* VNP20009 and their mutants were routinely cultured in LB broth. When needed, LB broth was supplemented with chloramphenicol (17 µg/mL), kanamycin (50 µg/mL) and/or carbenicillin (200 µg/mL). For *S. enterica* mutants lacking *asd* and *murI* genes, 250 µg/mL of DAP and D-glutamate were supplemented in the medium. For mass spectrometry analyses, strains were grown in M9 medium supplemented with 0.4% glucose, 0.1% CAS amino acids along with DAP, D-glutamate and antibiotics. For all experiments the culturing conditions were 37℃ with shaking.

### Genetic manipulation of *S. enterica*

For chromosomal gene knockout in *S. enterica* ATCC 14028s and *S. enterica* VNP20009, bacteria were first rendered ampicillin-resistant through transformation with plasmid p101-Amp containing an ampicillin-resistance cassette and a temperature-sensitive derivative of pSC101 replication origin. This plasmid was constructed by deleting λ red recombination genes from pKD46 plasmid^50^. Two PCR reactions were then used to amplify 0.8∼1 kb DNA fragments surrounding the sequence to be deleted in the *S. enterica* chromosome. The two homologous recombination arms were fused through a third PCR reaction, then ligated to a suicide plasmid using the In-Fusion® HD Cloning kit (Clontech). The suicide plasmid contained R6K replication origin, RP4-oriT, *sacB* as a counter selection marker, and a kanamycin-resistance cassette. The ligated suicide plasmid was transformed into *Escherichia coli* S17 λpir, then transferred to *S. enterica/* p101-Amp through conjugation. The resulting *S. enterica* merodiploid conjugants were plated on LB agar plates containing kanamycin at 50 ng/µl and carbenicillin at 200 ng/µl. One *S. enterica* merodiploid mutant was then selected and re-streaked on LSW-Sucrose agar plate ^51^. The composition of the LSW agar media was tryptone 10 g/l, yeast extract 5 g/l, glycerol 5 ml/l, NaCl 0.4 g/l, sucrose 100 g/l, and agar 20 g/l. The plates were supplemented with kanamycin at 50 ng/µl and either D-glutamate at 250 µg/ml (for *murI* knockouts) and/or DAP at 250 µg/ml (for *asd* knockouts). Screening for knockout mutants was done through PCR. The correct knockouts were selected, re-streaked, and confirmed for the loss of the conjugated plasmid through DNA sequencing and its inability to grow in the presence of kanamycin. To cure the knockouts from the ampicillin-resistance plasmid, they were cultured at 40 °C and screened for the loss of ampicillin resistance. For chromosomal knock-in of *kynU* gene cassette in *S. enterica* AD95, the same procedure was used; however, there was no need to transform it in advance with an antibiotic-resistant plasmid as this strain is both chloramphenicol and kanamycin resistant. Instead, the suicide plasmid harbored a carbenicillin-resistant cassette allowing for plating the merodiploid resulting from conjugation on LB plates supplemented with carbenicillin, kanamycin, and chloramphenicol. The kynU cassette was inserted within the previously deleted *asd* locus.

### Assessing growth of *S. enterica* mutants

*S. enterica* strains were cultured in LB broth for 24 h, followed by centrifugation and washing the pellets twice in M9 media. 2 µl of these suspensions were added to 200 µl of M9 media supplemented with glucose at 0.4% and CAS amino acids at 0.1% inside 96-well plates. M9 media were supplemented with chloramphenicol and kanamycin when appropriate. The plates were incubated at 37 °C with shaking within a Tecan Infinite microplate reader. OD_600_ was recorded at 30-minute intervals over 24 h period.

### Cell lines

Mouse ovarian epithelial cancer cell line KPCA.A^52^ was a gift from R. A. Weinberg at Whitehead Institute for Biomedical Research to the Reizes lab. KPCA.A was cultured in DMEM supplemented with 4% heat-inactivated fetal bovine serum (FBS), 1% insulin-transferrin-selenium (ITS-G, 41400045, Thermo Fisher Scientific), epidermal growth factor (2 ng/mL) and 100 U ml^−1^ penicillin–streptomycin. B16-F10 mouse melanoma cells were grown in DMEM containing 10% FBS and 100 U ml^−1^ penicillin–streptomycin. All cell lines were cultured at 37 °C with a humidified atmosphere and 5% CO_2_.

### Mice tumor experiments

All animal experiments were approved by the Institutional Animal Care and Use Committee of Cleveland Clinic Animal Care and Use Committee (IACUC No. 3279). To implant tumors in mice, respective cancer cells grown in vitro were collected, washed, and resuspended in 200 µL of PBS. To implant KPCA.A tumor, 10×10^6^ cells were injected subcutaneously into the right flank region of female C57BL/6J mice. For B16-F10 tumor model, 5×10^5^ B16-F10 cells were injected into the right flank region of both male and female C57BL/6J mice. Tumor volume was measured using calipers, and mice were assigned randomly to treatment groups once the tumor was of measurable size. For treatment, bacteria were cultured overnight as described before, washed two times using PBS and resuspended in PBS. Unless indicated otherwise, 200 µL of bacterial suspension (Equivalent to 1×10^7^ CFU) was injected intravenously through the retroorbital route. For the combination therapy, 5 µg of mouse IFNγ (AF-315-05-250UG, Thermo Fisher Scientific) in PBS was injected intratumorally. Bacterial and IFNγ administrations were followed as illustrated in the experimental schemes.

### Mass spectrometry

LC-MS/MS analyses for bacterial cultures and tumor samples were performed as described previously^33^. Briefly, the in vitro cultures were diluted as appropriate while tumors were mixed with 3 volumes of MQ water and incubated at 95℃ for 5 minutes. The samples were homogenized using a bead tissue homogenizer (MM400, Retsch). All samples, both in vitro and in vivo, were filtered through a 3-kDa molecular weight cutoff membrane filter (UFC5003BK, Amicon). The filtrate was mixed with 1/10 volume of 100 μM internal standard mix containing [^2^H_4_]-l-kynurenine (D-8026, CDN isotopes), [^2^H_5_]-l-tryptophan (D-1522, CDN isotopes) and [^2^H_5_]-kynurenic acid (D-439, CDN isotopes). 1 μl of the resulting filtrate mix was injected into LC-MS/MS spectrometer (API 5000 or API 5500 Mass spectrometer, Sciex) through a Shimadzu autosampler. A C18 column (Prodigy, 150 by 2 mm, 5 µm, 00F-3300-B0, Phenomenex) was used to resolve the metabolites using a gradient generated from solvent A, 0.2% formic acid in water and solvent B, 0.2% formic acid in methanol. A series of standard solutions containing kynurenine, tryptophan, kynurenic acid, and anthranilic acid at varying concentrations were prepared and mixed with the internal standard mixture at the same volume ratio as the study samples to generate calibration curves for each metabolite. Mass spectrometry (MS) parameters were optimized individually for each analyte and its corresponding internal standard.

### ELISA

Tumor samples were snap frozen immediately after dissection and stored at -80℃. Tumor samples were ground in liquid nitrogen and protein was extracted using T-PER tissue protein extraction reagent (78510, Thermo Scientific) containing protease inhibitor cocktail (11836170001, Roche) and phenylmethyl sulfonyl fluoride (7110, Sigma-Aldrich). Protein was estimated by Pierce Bicinchoninic Acid protein assay kit (23225, Thermo Scientific). 250 µg of total protein from KPCA.A tumors and 500 µg of total protein from B16-F10 tumors were used for cytokine analysis. Cytokines were detected by V-PLEX Proinflammatory Panel 1 Mouse Kit (K15048D, MSD). The analysis was done on a MESO QuickPlex SQ 120 instrument as per manufacturer’s instructions.

### Tumor dissociation and flow cytometry

Tumor samples were stored immediately after dissection at 4℃ in MACS Tissue Storage Solution (130-100-008, Miltenyi Biotech) overnight. The next day, tumor tissues were washed in PBS, minced into small pieces, and dissociated in 3 mL of DMEM containing collagenase (C5138, Sigma) 1 mg/mL and DNase I (11284932001, Roche) 0.1 mg/mL. The homogenate was pipetted up and down and incubated at 37 ℃ with shaking for 30 minutes. After incubation, the homogenate was again pipetted up and down gently, filtered through a 40 µm cell strainer (07000222, Greiner Bio-One), and centrifuged at 300×g for 7 minutes. RBCs lysis was performed using RBC lysis buffer (11814389001, Roche). Cells were washed using staining buffer (PBS containing 0.5% BSA). Cell counting was performed by TC20 Automated Cell Counter (Bio-Rad, USA), and 4 × 10^6^ cells were used for flow cytometry staining. Fc receptors were blocked by 15-min incubation with anti-mouse CD16/CD32 antibody (BDB553142, BD Biosciences) following which the cells were stained using the live/dead dye (Live-or-Dye™ 615/740, 32015 Biotium). The cells were washed and incubated with fluorochrome-conjugated antibodies for 30 min. The cells were then fixed overnight using ThermoFisher fixation/permeabilization buffer (00-5521-00 or 00-8222-49, ThermoFisher). Cells were fixed by addition of 200 μL of fixative per well. Following fixation, cells were spun down and washed in staining buffer twice. For intracellular staining, the cells were permeabilized by resuspending in 200 μL of permeabilization buffer (00-8333-56, ThermoFisher). The list of antibodies used is provided in Table S2. All incubations were performed on ice. Cells were finally washed and resuspended in the staining buffer before being analyzed by Sony ID7000 spectral cell analyzer. Fluorescent minus one controls were used to set gates whenever clear clustering of cells was not observed. The analysis was performed by FlowJo v10.10. FlowAI (flow rate only) plugin ^53^ was used to clean the data and DownSample plugin within FlowJo software was used to normalize the cell number before further downstream analysis. The flow cytometry gating strategy is shown in Figure S5.

### In vitro restimulation of T cells for cytokine analysis by flow cytometry

Following tumor dissociation, 4×10^6^ cells were incubated in 24 well plates containing 500 µL DMEM supplemented with 0.05 µg/mL phorbol myristate acetate (P1585, Sigma), 0.5 µg/mL ionomycin (I0634, Sigma) and 1X Brefeldin A (420601, Biolegend) for 3.5 h at 37°C in a humidified incubator supplemented with 5% CO_2_. Following incubation, the cells were washed twice using the staining buffer and proceeded with flow cytometry staining as explained above. Flow cytometry analysis was performed by Sony ID7000 spectral cell analyzer, and data were analysed by FlowJo v10.10.

### Bulk RNA sequencing and analysis

Library preparation, sequencing and DE analysis were performed by GENEWIZ® NGS Services from Azenta Life Sciences (South Plainfield, NJ, USA). Total RNA was extracted from five mice in each treatment group using Qiagen RNeasy Plus Mini kit (Qiagen, Hilden, Germany) following manufacturer’s instructions. RNA samples were quantified using Qubit 4.0 Fluorometer (ThermoFisher Scientific, Waltham, MA, USA) and RNA integrity was checked using 4200 TapeStation (Agilent Technologies, Palo Alto, CA, USA). rRNA depletion was done using QIAGEN FastSelect rRNA HMR Kit (Qiagen, Hilden, Germany). RNA sequencing library was prepared using NEBNext Ultra II RNA Library Preparation Kit following the manufacturer’s recommendations (NEB, Ipswich, MA, USA). The samples were sequenced on a NovaSeq platform using 2×150 bp paired end configuration. Image analysis and base calling were conducted by NovaSeq Control Software (NCS). Raw sequence data (bcl files) generated from Illumina NovaSeq was converted into fastq files and de-multiplexed using Illumina bcl2fastq 2.20 software. One mismatch was allowed for index sequence identification. Adapter sequences and poor-quality reads were trimmed using Trimmomatic v.0.36. The trimmed reads were mapped to the reference genome (GRCm38) using the STAR aligner v.2.5.2b. Unique gene hit counts were calculated by using feature Counts from the Subread package v.1.5.2. Only unique reads that fell within exon regions were counted. DESeq2 was used for differential expression (DE) analysis. The Wald test was used to generate p values and log2 fold changes. All genes that were altered significantly (p values < 0.05) following DE analysis, were subjected to pathway enrichment analysis using Ingenuity Pathway Analysis (IPA) software (Qiagen, Hilden, Germany). Heatmaps were generated by Morpheus (https://software.broadinstitute.org/morpheus) and IPA. Genes selected to generate heatmaps are those which showed significant (p values < 0.05) differential expression in at least in one of the datasets.

### Statistical analysis

One-way ANOVA was used to compare different groups followed by Tukey post-hoc test when the samples are normally distributed. Kruskal-Wallis (for un-matched groups) and Friedman tests (for matched groups) were used for statistical analyses of data which were not normally distributed. Student t-test, and Mann-Whitney U test were used for pair-wise comparisons. Analysis was performed using GraphPad Prism 11. For animal experiments, we used only female mice for KPCA.A ovarian tumor model while for B16-F10 melanoma model, both male and female mice were used. Mice were assigned randomly to different groups. Laboratory personnel were not blinded during the animal experiments to avoid cross-contamination between groups. Each point on the graphs represents an individual mouse.

## Acknowledgments

Part of the illustrations shown in the figures were created using BioRender.com. We thank Apryl Helmick and Bridget Besse in the Lerner Research Institute Imaging core. We thank Kimberly Peterson in the Lerner Research Institute Laboratory Diagnostic Core for her help with the analyses of cytokines level in mice tumor samples. We thank Kewal Asosingh and the rest of the team in the Cleveland Clinic Research Flow Cytometry Core Facility. The Sony ID7000 is supported by NIH grant S10OD025207-01A1. The International Society for Advancement of Cytometry (ISAC) recognizes Cleveland Clinic Research Flow Cytometry Core for exceeding the excellence threshold outlined in the society’s best practices for SRL operations. NIH R35 GM156762 grant supported the bacterial genetic engineering studies. We are grateful for the support.

## Author contributions

MD conceived and initiated the study and designed the experiments. MD pursued funding. MD wrote the initial manuscript draft. MD and KKV edited the manuscript. MD and AS performed the bacterial genetic engineering experiments. KKV and RB performed the tissue culture. MD, KKV, AMW and ZW performed the LC-MS/MS analyses. KKV performed flow cytometry experiments and analysis. KKV performed pathway enrichment analysis. KA, PA and NZ contributed to the flow cytometry protocols and analyses. MD, KKV, AS, TX, MA, and RB performed the mice experiments. MD supervised the work. All authors provided critical input on data analyses. All authors agreed to the final version of the manuscript.

## Data and materials availability

All data are available in the main text or the supplementary materials. Genetic constructs made during this study can be reasonably requested by other investigators from Dr. Mohammed Dwidar after submission of appropriate institutional Materials Transfer Agreement between Cleveland Clinic Foundation and their research institutes.

## References

1. Amobi-McCloud, A., Muthuswamy, R., Battaglia, S., Yu, H., Liu, T., Wang, J., Putluri, V., Singh, P.K., Qian, F., Huang, R.Y., et al. (2021). IDO1 Expression in Ovarian Cancer Induces PD-1 in T Cells via Aryl Hydrocarbon Receptor Activation. Front Immunol 12, 678999. 10.3389/fimmu.2021.678999.

2. de Jong, R.A., Nijman, H.W., Boezen, H.M., Volmer, M., Ten Hoor, K.A., Krijnen, J., van der Zee, A.G., Hollema, H., and Kema, I.P. (2011). Serum tryptophan and kynurenine concentrations as parameters for indoleamine 2,3-dioxygenase activity in patients with endometrial, ovarian, and vulvar cancer. Int J Gynecol Cancer 21, 1320–1327. 10.1097/IGC.0b013e31822017fb.

3. Smith, L.P., Bitler, B.G., Richer, J.K., and Christenson, J.L. (2019). Tryptophan catabolism in epithelial ovarian carcinoma. Trends Cancer Res 14, 1–9.

4. Adams, S., Teo, C., McDonald, K.L., Zinger, A., Bustamante, S., Lim, C.K., Sundaram, G., Braidy, N., Brew, B.J., and Guillemin, G.J. (2014). Involvement of the kynurenine pathway in human glioma pathophysiology. PLoS One 9, e112945. 10.1371/journal.pone.0112945.

5. Riess, C., Schneider, B., Kehnscherper, H., Gesche, J., Irmscher, N., Shokraie, F., Classen, C.F., Wirthgen, E., Domanska, G., Zimpfer, A., et al. (2020). Activation of the Kynurenine Pathway in Human Malignancies Can Be Suppressed by the Cyclin-Dependent Kinase Inhibitor Dinaciclib. Front Immunol 11, 55. 10.3389/fimmu.2020.00055.

6. Zimmer, P., Schmidt, M.E., Prentzell, M.T., Berdel, B., Wiskemann, J., Kellner, K.H., Debus, J., Ulrich, C., Opitz, C.A., and Steindorf, K. (2019). Resistance Exercise Reduces Kynurenine Pathway Metabolites in Breast Cancer Patients Undergoing Radiotherapy. Front Oncol 9, 962. 10.3389/fonc.2019.00962.

7. Sakurai, K., Amano, S., Enomoto, K., Kashio, M., Saito, Y., Sakamoto, A., Matsuo, S., Suzuki, M., Kitajima, A., Hirano, T., and Negishi, N. (2005). [Study of indoleamine 2,3-dioxygenase expression in patients with breast cancer]. Gan To Kagaku Ryoho 32, 1546–1549.

8. Heng, B., Bilgin, A.A., Lovejoy, D.B., Tan, V.X., Milioli, H.H., Gluch, L., Bustamante, S., Sabaretnam, T., Moscato, P., Lim, C.K., and Guillemin, G.J. (2020). Differential kynurenine pathway metabolism in highly metastatic aggressive breast cancer subtypes: beyond IDO1-induced immunosuppression. Breast Cancer Res 22, 113. 10.1186/s13058-020-01351-1.

9. Heng, B., Lim, C.K., Lovejoy, D.B., Bessede, A., Gluch, L., and Guillemin, G.J. (2016). Understanding the role of the kynurenine pathway in human breast cancer immunobiology. Oncotarget 7, 6506–6520. 10.18632/oncotarget.6467.

10. Venkateswaran, N., Lafita-Navarro, M.C., Hao, Y.H., Kilgore, J.A., Perez-Castro, L., Braverman, J., Borenstein-Auerbach, N., Kim, M., Lesner, N.P., Mishra, P., et al. (2019). MYC promotes tryptophan uptake and metabolism by the kynurenine pathway in colon cancer. Genes Dev 33, 1236–1251. 10.1101/gad.327056.119.

11. Crotti, S., Fraccaro, A., Bedin, C., Bertazzo, A., Di Marco, V., Pucciarelli, S., and Agostini, M. (2020). Tryptophan Catabolism and Response to Therapy in Locally Advanced Rectal Cancer (LARC) Patients. Front Oncol 10, 583228. 10.3389/fonc.2020.583228.

12. Sun, X.Z., Zhao, D.Y., Zhou, Y.C., Wang, Q.Q., Qin, G., and Yao, S.K. (2020). Alteration of fecal tryptophan metabolism correlates with shifted microbiota and may be involved in pathogenesis of colorectal cancer. World J Gastroenterol 26, 7173–7190. 10.3748/wjg.v26.i45.7173.

13. Lin, D.J., Ng, J.C.K., Huang, L., Robinson, M., O’Hara, J., Wilson, J.A., and Mellor, A.L. (2021). The immunotherapeutic role of indoleamine 2,3-dioxygenase in head and neck squamous cell carcinoma: A systematic review. Clin Otolaryngol 46, 919–934. 10.1111/coa.13794.

14. Uyttenhove, C., Pilotte, L., Theate, I., Stroobant, V., Colau, D., Parmentier, N., Boon, T., and Van den Eynde, B.J. (2003). Evidence for a tumoral immune resistance mechanism based on tryptophan degradation by indoleamine 2,3-dioxygenase. Nat Med 9, 1269–1274. 10.1038/nm934.

15. Puccetti, P., Fallarino, F., Italiano, A., Soubeyran, I., MacGrogan, G., Debled, M., Velasco, V., Bodet, D., Eimer, S., Veldhoen, M., et al. (2015). Accumulation of an Endogenous Tryptophan-Derived Metabolite in Colorectal and Breast Cancers. Plos One 10. ARTN e0122046 10.1371/journal.pone.0122046.

16. Nguyen, N.T., Nakahama, T., Le, D.H., Van Son, L., Chu, H.H., and Kishimoto, T. (2014). Aryl hydrocarbon receptor and kynurenine: recent advances in autoimmune disease research. Front Immunol 5, 551. 10.3389/fimmu.2014.00551.

17. Munn, D.H., and Mellor, A.L. (2016). IDO in the Tumor Microenvironment: Inflammation, Counter-Regulation, and Tolerance. Trends Immunol 37, 193–207. 10.1016/j.it.2016.01.002.

18. Triplett, T.A., Garrison, K.C., Marshall, N., Donkor, M., Blazeck, J., Lamb, C., Qerqez, A., Dekker, J.D., Tanno, Y., Lu, W.C., et al. (2018). Reversal of indoleamine 2,3-dioxygenase-mediated cancer immune suppression by systemic kynurenine depletion with a therapeutic enzyme. Nat Biotechnol 36, 758–764. 10.1038/nbt.4180.

19. Holmgaard, R.B., Zamarin, D., Li, Y., Gasmi, B., Munn, D.H., Allison, J.P., Merghoub, T., and Wolchok, J.D. (2015). Tumor-Expressed IDO Recruits and Activates MDSCs in a Treg-Dependent Manner. Cell Rep 13, 412–424. 10.1016/j.celrep.2015.08.077.

20. Campesato, L.F., Budhu, S., Tchaicha, J., Weng, C.-H., Gigoux, M., Cohen, I.J., Redmond, D., Mangarin, L., Pourpe, S., Liu, C., et al. (2020). Blockade of the AHR restricts a Treg-macrophage suppressive axis induced by L-Kynurenine. Nat Commun 11, 4011. 10.1038/s41467-020-17750-z.

21. Triplett, T.A., Garrison, K.C., Marshall, N., Donkor, M., Blazeck, J., Lamb, C., Qerqez, A., Dekker, J.D., Tanno, Y., Lu, W.-C., et al. (2018). Reversal of indoleamine 2,3-dioxygenase– mediated cancer immune suppression by systemic kynurenine depletion with a therapeutic enzyme. Nat Biotechnol 36, 758–764. 10.1038/nbt.4180.

22. Odunsi, K., Qian, F., Lugade, A.A., Yu, H., Geller, M.A., Fling, S.P., Kaiser, J.C., Lacroix, A.M., D’Amico, L., Ramchurren, N., et al. (2022). Metabolic adaptation of ovarian tumors in patients treated with an IDO1 inhibitor constrains antitumor immune responses. Sci Transl Med 14, eabg8402. 10.1126/scitranslmed.abg8402.

23. Knoten, C.A., Hudson, L.L., Coleman, J.P., Farrow, J.M., 3rd, and Pesci, E.C. (2011). KynR, a Lrp/AsnC-type transcriptional regulator, directly controls the kynurenine pathway in Pseudomonas aeruginosa. J Bacteriol 193, 6567–6575. 10.1128/JB.05803-11.

24. Bortolotti, P., Hennart, B., Thieffry, C., Jausions, G., Faure, E., Grandjean, T., Thepaut, M., Dessein, R., Allorge, D., Guery, B.P., et al. (2016). Tryptophan catabolism in Pseudomonas aeruginosa and potential for inter-kingdom relationship. BMC Microbiol 16, 137. 10.1186/s12866-016-0756-x.

25. Kurnasov, O., Jablonski, L., Polanuyer, B., Dorrestein, P., Begley, T., and Osterman, A. (2003). Aerobic tryptophan degradation pathway in bacteria: novel kynurenine formamidase. FEMS Microbiol Lett 227, 219–227. 10.1016/S0378-1097(03)00684-0.

26. Farrow, J.M., 3rd, and Pesci, E.C. (2007). Two distinct pathways supply anthranilate as a precursor of the Pseudomonas quinolone signal. J Bacteriol 189, 3425–3433. 10.1128/JB.00209-07.

27. Chein, T., Doshi, A., and Danino, T. (2017). Advances in bacterial cancer therapies using synthetic biology. Current Opinion in Systems Biology 5, 8.

28. Daschner, P.J., Rasooly, A., and White, J.D. (2019). Bugs as Cancer Drugs: Challenges and Opportunities. Mol Cell Biol 39. 10.1128/MCB.00206-19.

29. Salicrup, L.A., Ossandon, M., Prickril, B., and Rasooly, A. (2020). Bugs as Drugs, potential self-regenerated innovative cancer therapeutics approach for global health. J Glob Health 10, 010311. 10.7189/jogh.10.010311.

30. Mehta, N., Lyon, J.G., Patil, K., Mokarram, N., Kim, C., and Bellamkonda, R.V. (2017). Bacterial Carriers for Glioblastoma Therapy. Mol Ther Oncolytics 4, 1–17. 10.1016/j.omto.2016.12.003.

31. Duong, M.T., Qin, Y., You, S.H., and Min, J.J. (2019). Bacteria-cancer interactions: bacteria-based cancer therapy. Exp Mol Med 51, 1–15. 10.1038/s12276-019-0297-0.

32. Varsha, K.K., and Dwidar, M. (2026). Engineering Salmonella as an immune-metabolic modulator of the tumor microenvironment. Trends Biotechnol. 10.1016/j.tibtech.2026.01.010.

33. Santos, A., Wang, Z., Bharti, R., Dey, G., Sangwan, N., Baldwin, W., Zalavadia, A., Myers, A., Huffman, O.G., Lathia, J.D., et al. (2025). Leveraging dysregulated tumor metabolism for targeting anticancer bacteria. Sci Adv 11, eads1630. 10.1126/sciadv.ads1630.

34. Pawelek, J.M., Low, K.B., and Bermudes, D. (1997). Tumor-targeted Salmonella as a novel anticancer vector. Cancer Res 57, 4537–4544.

35. Low, K.B., Ittensohn, M., Le, T., Platt, J., Sodi, S., Amoss, M., Ash, O., Carmichael, E., Chakraborty, A., Fischer, J., et al. (1999). Lipid A mutant Salmonella with suppressed virulence and TNFalpha induction retain tumor-targeting in vivo. Nat Biotechnol 17, 37–41. 10.1038/5205.

36. Danino, T., Lo, J., Prindle, A., Hasty, J., and Bhatia, S.N. (2012). In Vivo Gene Expression Dynamics of Tumor-Targeted Bacteria. ACS Synth Biol 1, 465–470. 10.1021/sb3000639.

37. Toso, J.F., Gill, V.J., Hwu, P., Marincola, F.M., Restifo, N.P., Schwartzentruber, D.J., Sherry, R.M., Topalian, S.L., Yang, J.C., Stock, F., et al. (2002). Phase I study of the intravenous administration of attenuated *Salmonella* Typhimurium to patients with metastatic melanoma. J Clin Oncol 20, 142–152. 10.1200/JCO.2002.20.1.142.

38. Han, Q., Fang, J., and Li, J. (2001). Kynurenine aminotransferase and glutamine transaminase K of Escherichia coli: identity with aspartate aminotransferase. Biochem J 360, 617–623. 10.1042/0264-6021:3600617.

39. Jansen, R.S., Mandyoli, L., Hughes, R., Wakabayashi, S., Pinkham, J.T., Selbach, B., Guinn, K.M., Rubin, E.J., Sacchettini, J.C., and Rhee, K.Y. (2020). Aspartate aminotransferase Rv3722c governs aspartate-dependent nitrogen metabolism in Mycobacterium tuberculosis. Nat Commun 11, 1960. 10.1038/s41467-020-15876-8.

40. Doublet, P., van Heijenoort, J., Bohin, J.P., and Mengin-Lecreulx, D. (1993). The murI gene of Escherichia coli is an essential gene that encodes a glutamate racemase activity. J Bacteriol 175, 2970–2979. 10.1128/jb.175.10.2970-2979.1993.

41. Iyer, S., Zhang, S., Yucel, S., Horn, H., Smith, S.G., Reinhardt, F., Hoefsmit, E., Assatova, B., Casado, J., Meinsohn, M.C., et al. (2021). Genetically Defined Syngeneic Mouse Models of Ovarian Cancer as Tools for the Discovery of Combination Immunothrapy. Cancer Discovery 11, 384–407. 10.1158/2159-8290.Cd-20-0818.

42. Waters, C.M., and Bassler, B.L. (2005). Quorum sensing: cell-to-cell communication in bacteria. Annu Rev Cell Dev Biol 21, 319–346. 10.1146/annurev.cellbio.21.012704.131001.

43. Smith, D., Wang, J.H., Swatton, J.E., Davenport, P., Price, B., Mikkelsen, H., Stickland, H., Nishikawa, K., Gardiol, N., Spring, D.R., and Welch, M. (2006). Variations on a theme: diverse N-acyl homoserine lactone-mediated quorum sensing mechanisms in gram-negative bacteria. Sci Prog 89, 167–211. 10.3184/003685006783238335.

44. Murphy, B., Miyamoto, T., Manning, B.S., Mirji, G., Ugolini, A., Kannan, T., Hamada, K., Zhu, Y.P., Claiborne, D.T., Huang, L., et al. (2024). Myeloid activation clears ascites and reveals IL27-dependent regression of metastatic ovarian cancer. J Exp Med 221. 10.1084/jem.20231967.

45. Sun, L., Kees, T., Almeida, A.S., Liu, B., He, X.Y., Ng, D., Han, X., Spector, D.L., McNeish, I.A., Gimotty, P., et al. (2021). Activating a collaborative innate-adaptive immune response to control metastasis. Cancer Cell 39, 1361–1374 e1369. 10.1016/j.ccell.2021.08.005.

46. Opitz, C.A., Somarribas Patterson, L.F., Mohapatra, S.R., Dewi, D.L., Sadik, A., Platten, M., and Trump, S. (2020). The therapeutic potential of targeting tryptophan catabolism in cancer. Br J Cancer 122, 30–44. 10.1038/s41416-019-0664-6.

47. Wirthgen, E., Hoeflich, A., Rebl, A., and Gunther, J. (2017). Kynurenic Acid: The Janus-Faced Role of an Immunomodulatory Tryptophan Metabolite and Its Link to Pathological Conditions. Front Immunol 8, 1957. 10.3389/fimmu.2017.01957.

48. Jorgovanovic, D., Song, M., Wang, L., and Zhang, Y. (2020). Roles of IFN-gamma in tumor progression and regression: a review. Biomark Res 8, 49. 10.1186/s40364-020-00228-x.

49. Mojic, M., Takeda, K., and Hayakawa, Y. (2017). The Dark Side of IFN-gamma: Its Role in Promoting Cancer Immunoevasion. Int J Mol Sci 19. 10.3390/ijms19010089.

50. Datsenko, K.A., and Wanner, B.L. (2000). One-step inactivation of chromosomal genes in Escherichia coli K-12 using PCR products. Proc Natl Acad Sci U S A 97, 6640–6645. 10.1073/pnas.120163297.

51. Howery, K.E., and Rather, P.N. (2019). Allelic Exchange Mutagenesis in Proteus mirabilis. Methods Mol Biol 2021, 77–84. 10.1007/978-1-4939-9601-8_8.

52. Iyer, S., Zhang, S., Yucel, S., Horn, H., Smith, S.G., Reinhardt, F., Hoefsmit, E., Assatova, B., Casado, J., Meinsohn, M.C., et al. (2021). Genetically Defined Syngeneic Mouse Models of Ovarian Cancer as Tools for the Discovery of Combination Immunotherapy. Cancer Discov 11, 384–407. 10.1158/2159-8290.CD-20-0818.

53. Monaco, G., Chen, H., Poidinger, M., Chen, J., de Magalhaes, J.P., and Larbi, A. (2016). flowAI: automatic and interactive anomaly discerning tools for flow cytometry data. Bioinformatics 32, 2473–2480. 10.1093/bioinformatics/btw191.

