## Supplemental information file for "Metabolic engineering of *Salmonella enterica* for coupled kynurenine sensing and depletion enhances antitumor efficacy"

**Figure S1**

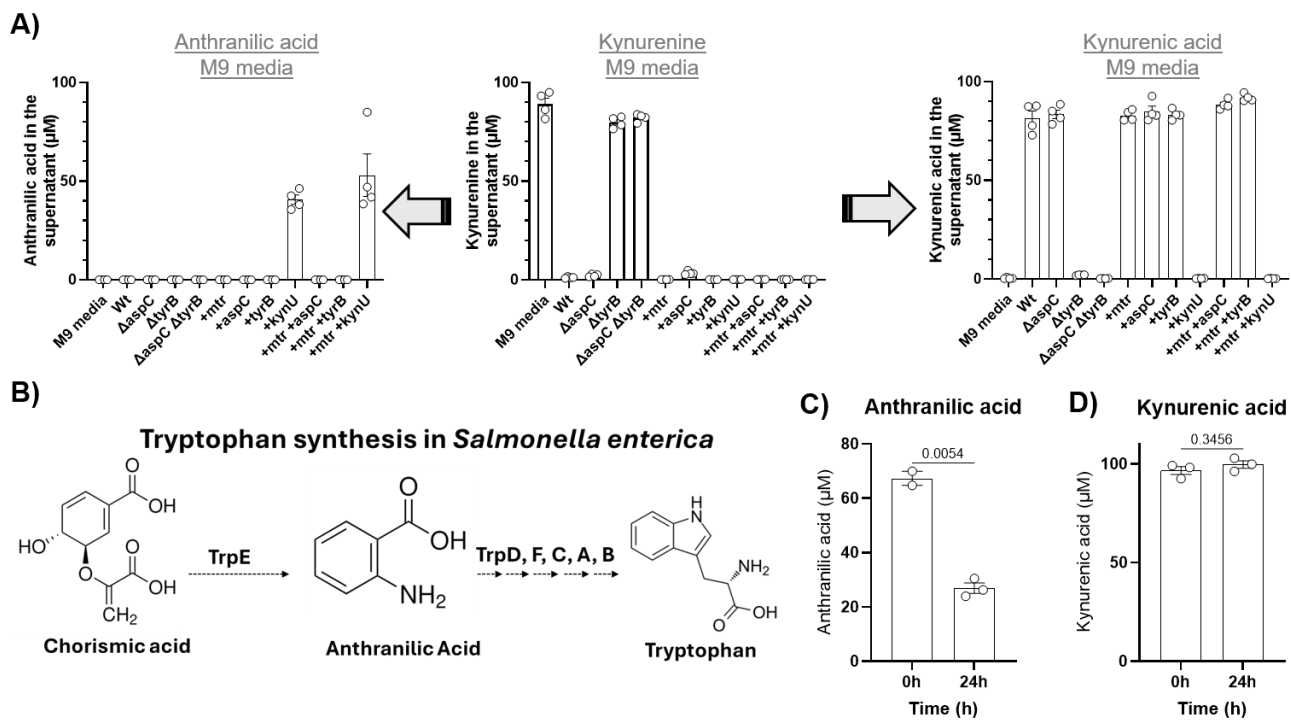

**Figure S1. Kynurenine metabolism by wildtype *S. enterica* and mutants in M9 media.** **A)** *S. enterica* and each of the respective mutants were cultured in M9 media supplemented with 0.4% glucose, 1% casamino acids and 100 μM kynurenine. Samples were analyzed after 24 h of incubation through LC-MS/MS. n=4. Plotted are means ± SE. **B)** Tryptophan biosynthetic pathway in Wild-type *S. enterica* **C)** Wildtype *S. enterica* was cultured in M9 media supplemented with 0.4% glucose, 1% casamino acids and either kynurenic acid or anthranilic acid for 24 hours. The spent media were then analyzed for the consumption of each of these two respective metabolites through LC-MS/MS. Plotted are means ± SE.

**Figure S2**

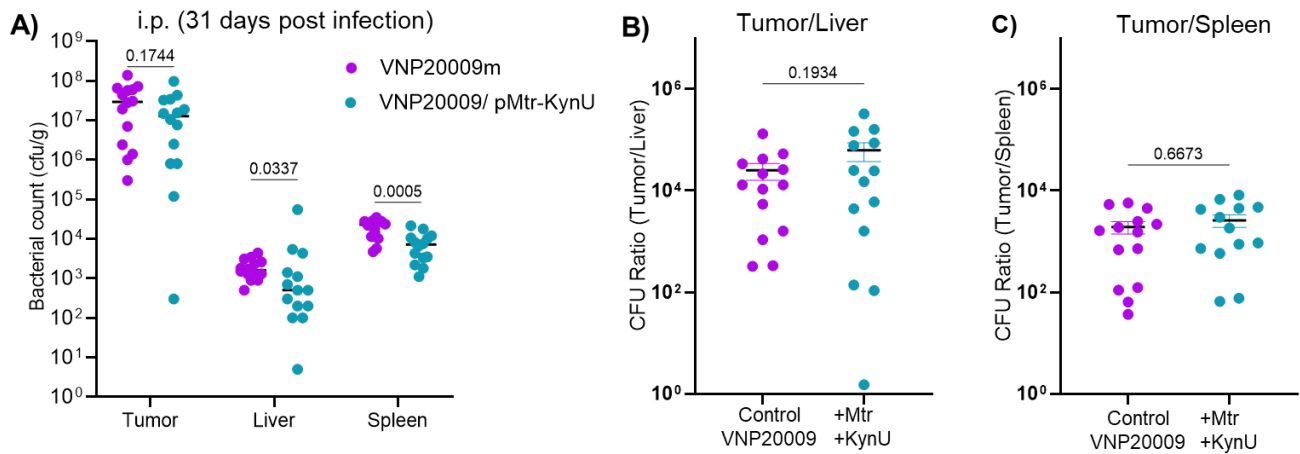

**Figure S2. CFU counting results for *S. enterica* VNP20009 and its kynurenine-degrading derivative (VMP20009/pMtr-KynU) for experiments shown in Figure 2.** KPCA.A tumors were developed subcutaneously in C57BL/6 mice. When the tumors were evident, VNP20009 mutants were intraperitoneally injected at a dose of $\sim 2 \times 10^6$ CFU weekly. The mice were euthanized 31 days after the tumor injection (when the tumors in the PBS group reached the endpoint). Data was combined from two separate experiments. Statistical analyses were done using Mann Whitney test. p-values are presented in the panels. Plotted are means ± SE.

**Figure S3**

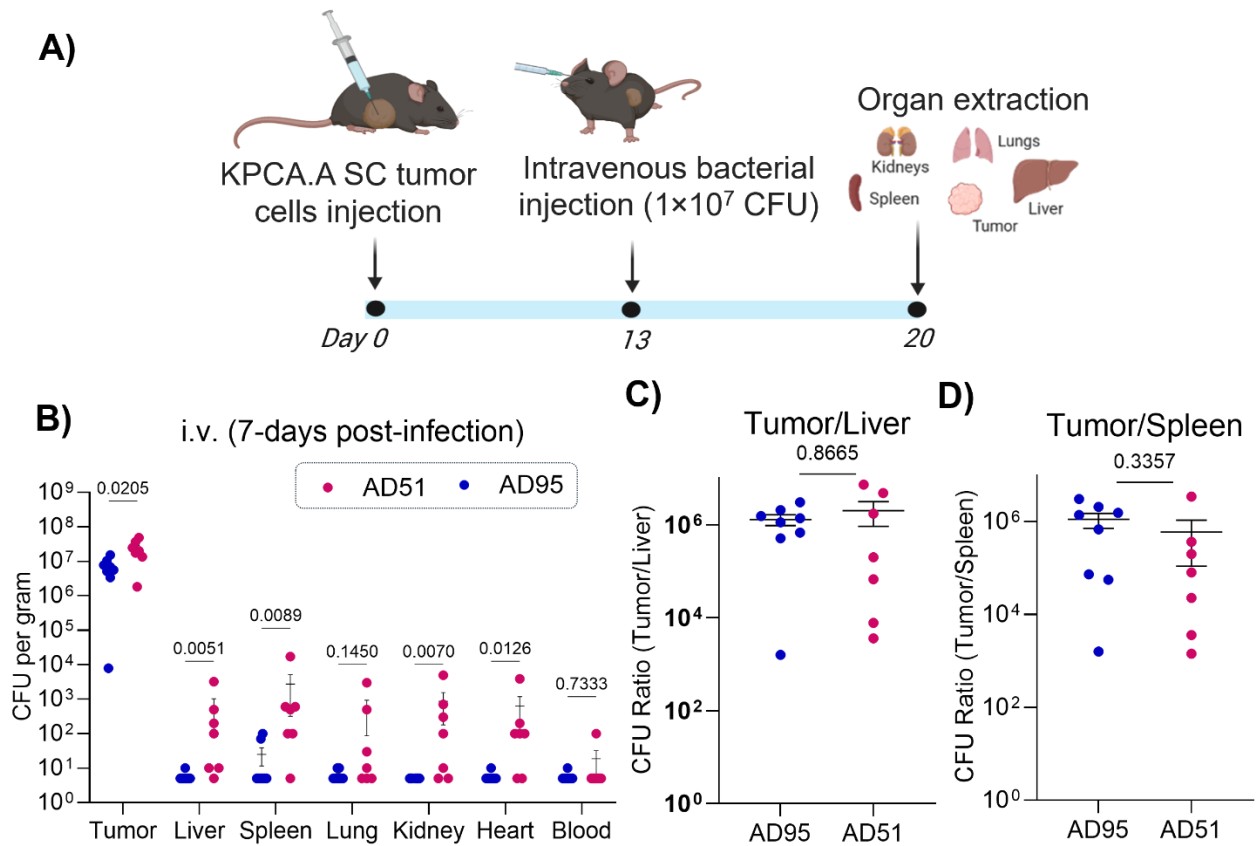

**Figure S3. Colonization by *S. enterica* AD95 and AD51 in SC KPCA.A tumors-bearing mice 7-days after intravenous injection.** **A)** Subcutaneous KPCA.A tumors were injected in C57BL/6 mice. When tumors reached a medium size, *Salmonella* mutants were intravenously injected at a dose of $\sim 1 \times 10^7$ CFU via retro-orbital route. **B-D)** Mice were euthanized 7 days later, and organs harvested for CFU counting ($n=7-8$). Each symbol represents results from an individual mouse. When no colonies were detected at the highest dilution, the number was stated as 5 CFU/g tissue which is half the limit of detection. Statistical analysis was done by Mann-Whitney test. p-values are presented in the panels. Plotted are means ± SE

Figure S4

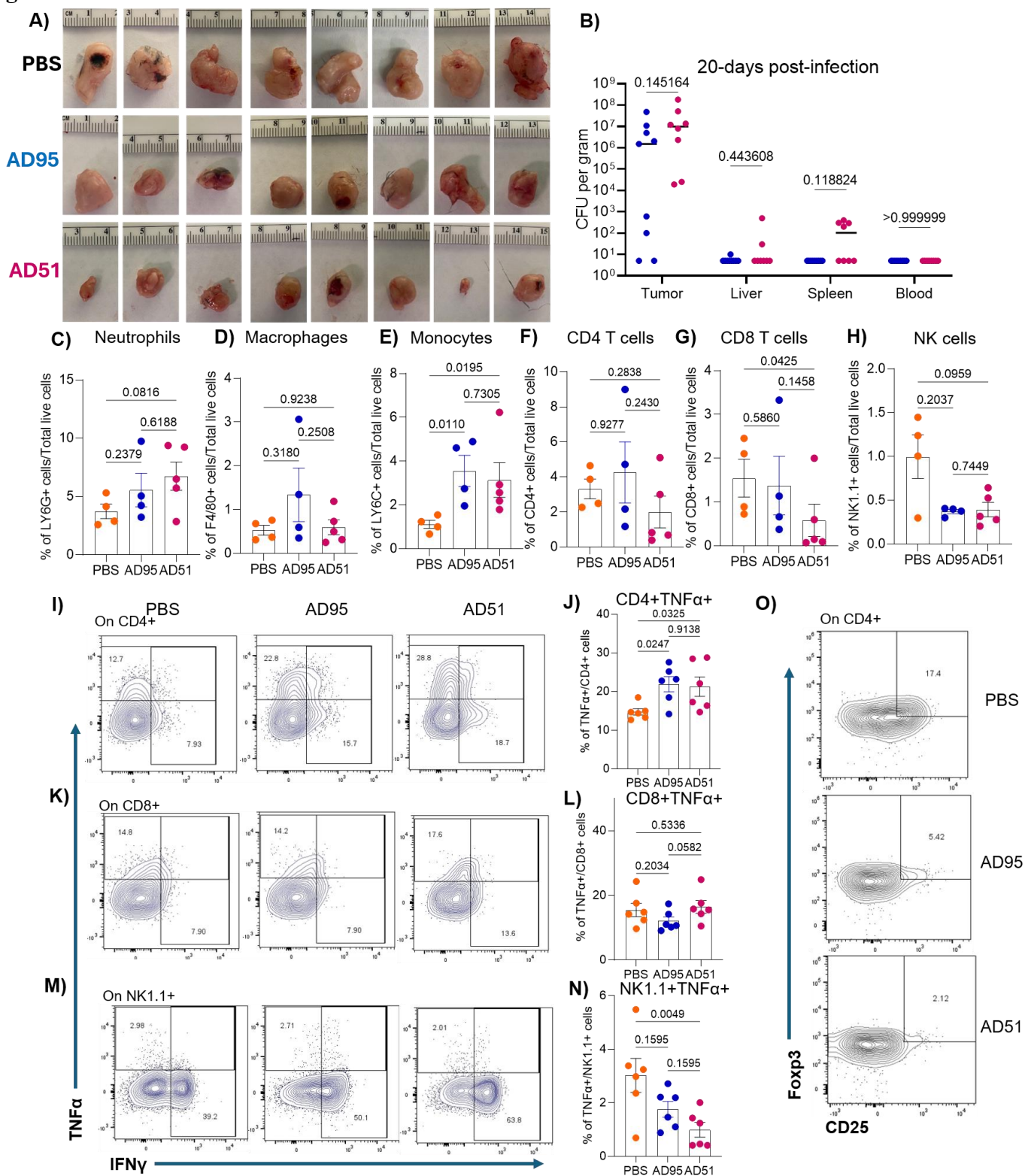

**Figure S4. Flow cytometry analysis demonstrated several immunostimulatory effects for AD51 compared to AD95 treatment. A)** Tumors collected from PBS and *Salmonella* treated mice. **B).** *Salmonella* tumor colonization at the end point. **C-O)** Flow cytometry analyses of the immune cells. **C-H)** Various immune cell populations within the tumor as percentage of total live cells of the tumor. **I-N)** TNFα production by CD4<sup>+</sup>, CD8<sup>+</sup> and NK cells. **O)** Flow cytometry plots for Treg data shown in Figure 4S. Statistical analyses were done using Kruskal-Wallis test. Plotted are means ± SE.

**Figure S5**

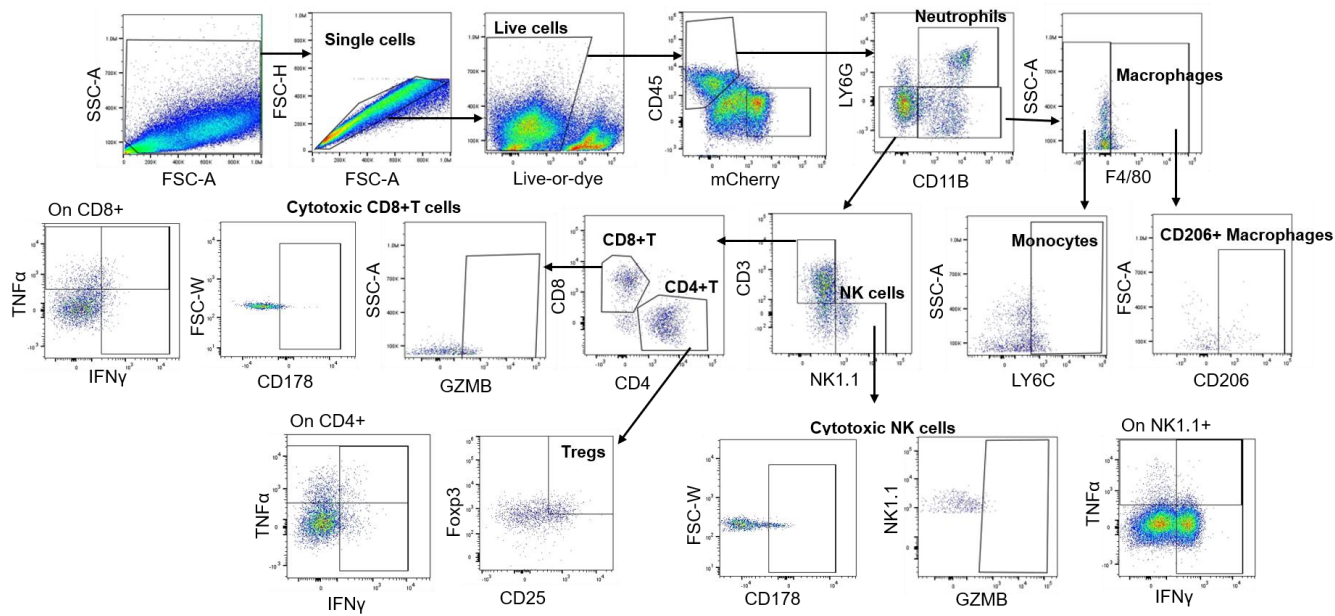

**Figure S5.** Gating strategy followed for analysis of flow cytometry data.

Figure S6

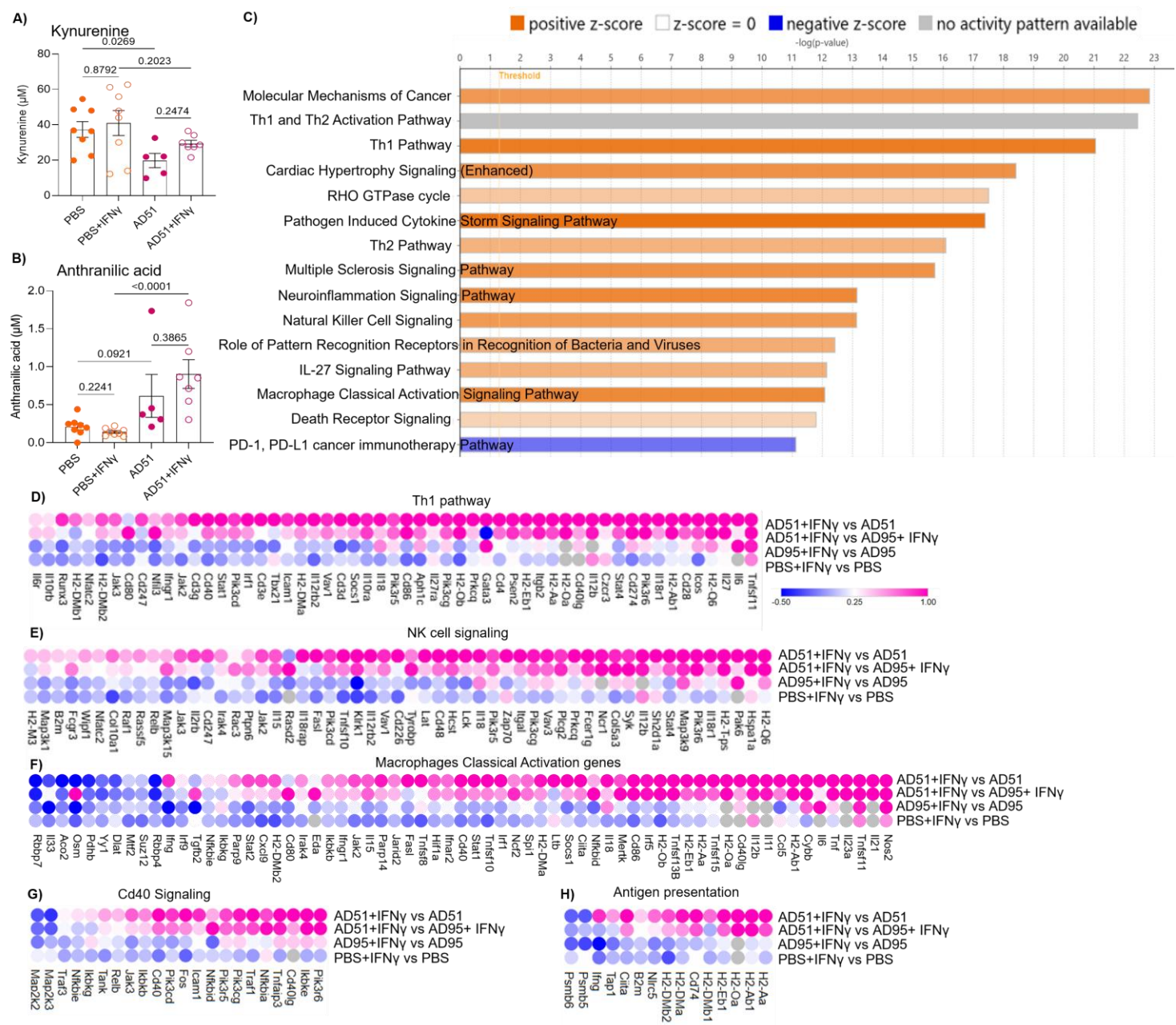

**Figure S6. Bulk transcriptomic analysis reveals broad immune pathway activation in KPCA.A tumors following AD51+IFNγ combination therapy. A) Tumor kynurenine and B) Tumor anthranilic acid. Statistical analyses were done using Kruskal-Wallis test. Plotted are means ± SE. C) Top 15 pathways altered in AD51+IFNγ group compared to AD51 treatment group. The displayed z-score predicts the activity state of that pathway. D-H) Heatmaps of expression level of genes associated with selected**

pathways including **D)** Th1 pathway, **E)** NK cell signaling, **F)** Macrophage classical activation, **G)** Cd40 signaling, and **H)** Antigen presentation.

**Figure S7**

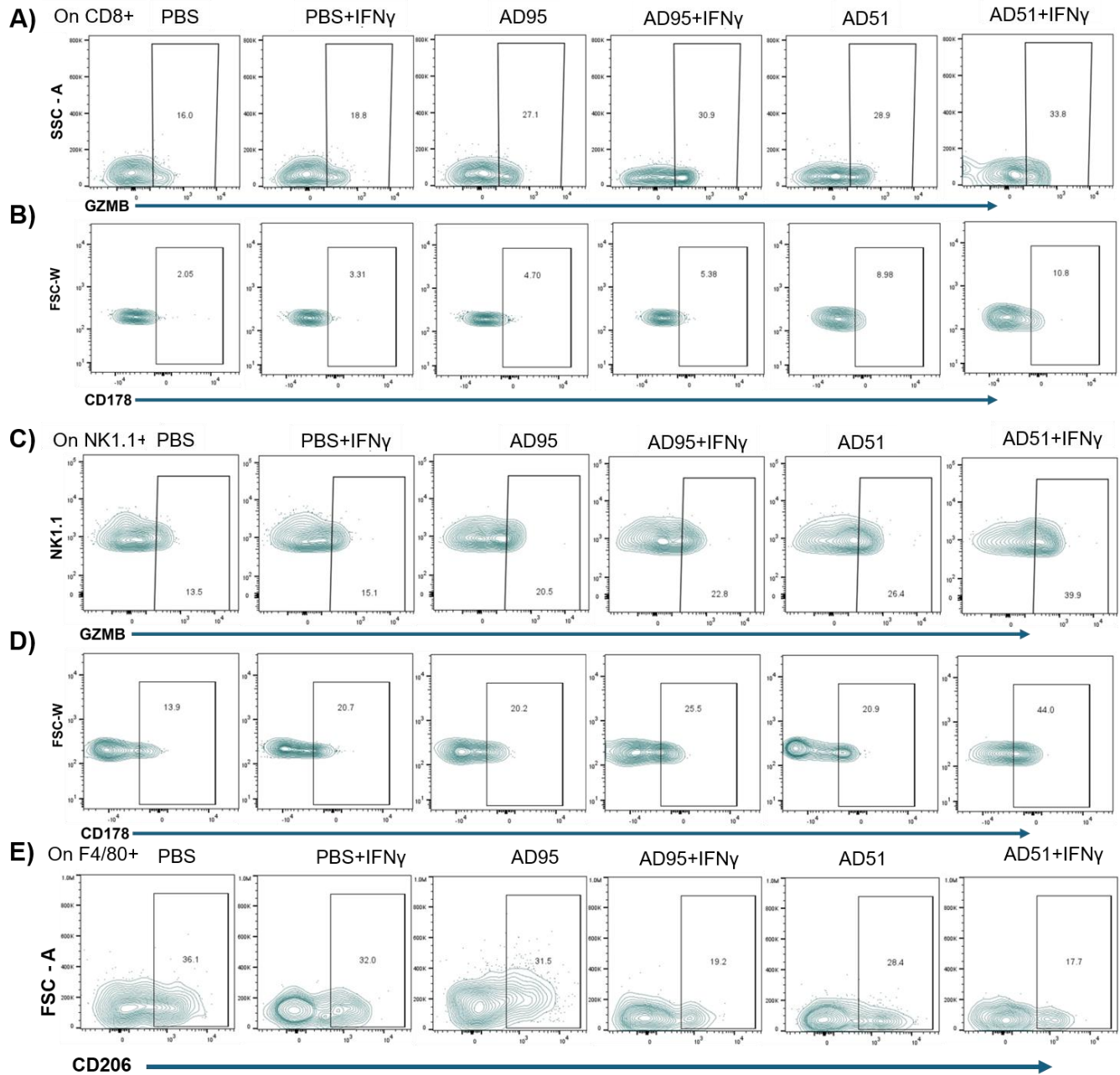

**Figure S7.** Flow cytometry analysis demonstrates enhanced immune activation in KPCA.A Tumors after AD51+IFNγ combination therapy **A)** GZMB+CD8+ T cells. **B)** CD178+CD8+ T cells. **C)** GZMB+ NK1.1+ cells. **D)** CD178+NK1.1+ cells. **E)** CD206+F4/80+ cells.
